# EVE is an open modular data analysis software for event-based localization microscopy

**DOI:** 10.1101/2024.08.09.607224

**Authors:** Laura M. Weber, Koen J.A. Martens, Clément Cabriel, Joel J. Gates, Manon Albecq, Fredrik Vermeulen, Katarina Hein, Ignacio Izeddin, Ulrike Endesfelder

## Abstract

Event-based sensors (EBS), or neuromorphic vision sensors, offer a novel approach to imaging by recording light intensity changes asynchronously, unlike conventional cameras that capture light over fixed exposure times. This capability results in high temporal resolution, reduced data redundancy, and a wide dynamic range. This makes EBS ideal for Single-Molecule Localization Microscopy (SMLM) as SMLM relies on the sequential imaging of sparse, blinking fluorescent emitters to achieve super-resolution. Recent studies have shown that EBS can effectively capture these emitters, achieving spatial resolution comparable to traditional cameras. However, existing analyses of event-based SMLM (eveSMLM) data have relied on converting event lists into image frames for conventional analysis, limiting the full potential of the technology.

To overcome this limitation, we developed EVE, a specialized software for analyzing eveSMLM data. EVE offers an integrated platform for detection, localization, and post-processing, with various algorithmic options tailored for the unique structure of eveSMLM data. EVE is user-friendly and features an open, modular infrastructure that supports ongoing development and optimization. EVE is the first dedicated tool for event-based SMLM, transforming the analysis process to fully utilize the spatiotemporal data generated by EBS. This allows researchers to explore the full potential of eveSMLM and encourages the development of new analytical methods and experimental improvements.

## Main text

Event-based sensors (EBS), also known as neuromorphic vision sensors, represent a promising development in imaging technology. Unlike conventional cameras such as electron multiplying charge-coupled devices (emCCDs) and scientific complementary metal–oxide–semiconductors (sCMOS) that integrate light intensities over a set exposure time, EBS record changes in incident light intensity by triggering events for individual pixels when passing predefined thresholds. This asynchronous event-readout offers several technological advantages, including high temporal resolution, greatly reduced data redundancy due to static signal suppression, and wide dynamic range due to its logarithmic response. This makes EBS ideally suited for recording single-molecule localization microscopy (SMLM) data, as this super-resolution technique relies on the sequential imaging of sparse, blinking fluorescent emitters.^1,2,3^ First work^4,5^ has shown that EBS are sensitive enough to record single emitters and to provide “eveSMLM” data with a spatial resolution similar to that of conventional cameras. Until now, however, analysis of eveSMLM data was performed by converting the event lists to image frames followed by traditional frame-based analysis. Thus, to date, the potential of the sensor for SMLM imaging, i.e. the achievable resolution or ability for multi-modal recordings, remains unexplored. We developed EVE to establish and integrate advanced eveSMLM data analysis and to further explore the scientific capacity of eveSMLM.

EVE is an integral detection, localization and post-processing software designed for eveSMLM experiments, offering several algorithmic options for all analysis steps (analysis methods implemented in EVE, Supplementary Note 1). The software is highly user-friendly due to its simple installation and graphical interface (user manual, Supplementary Note 2). Furthermore, EVE provides an open and modular infrastructure (developers instructions, Supplementary Note 3). This facilitates easy development and testing of future analysis routines, and optimization of experimental and hardware settings, promoting future advancements in eveSMLM analysis.

The EVE infrastructure is divided into three modules, each serving a distinct purpose in the analysis pipeline for eveSMLM data (Figure 1a). Event data is stored in lists, with each event having a (x,y) pixel coordinate and a microsecond-precise time stamp. Most sensors also include polarity information (p) indicating whether the intensity change is positive or negative. For those, analysis can be performed for each polarity separately or in combination. After loading the event data, the “Candidate Finding” module (Figure 1a, box 2) filters the raw input and groups candidates for events belonging to the same emitter into clusters. This identifies and extracts potential single-emitter signals. Besides the previously-used frame-based approach^4,5^, EVE offers new algorithms that directly analyze the event data by density-based clustering (Supplementary Note 1). A detailed preview helps the user make informed parameter decisions.

**Figure 1:**
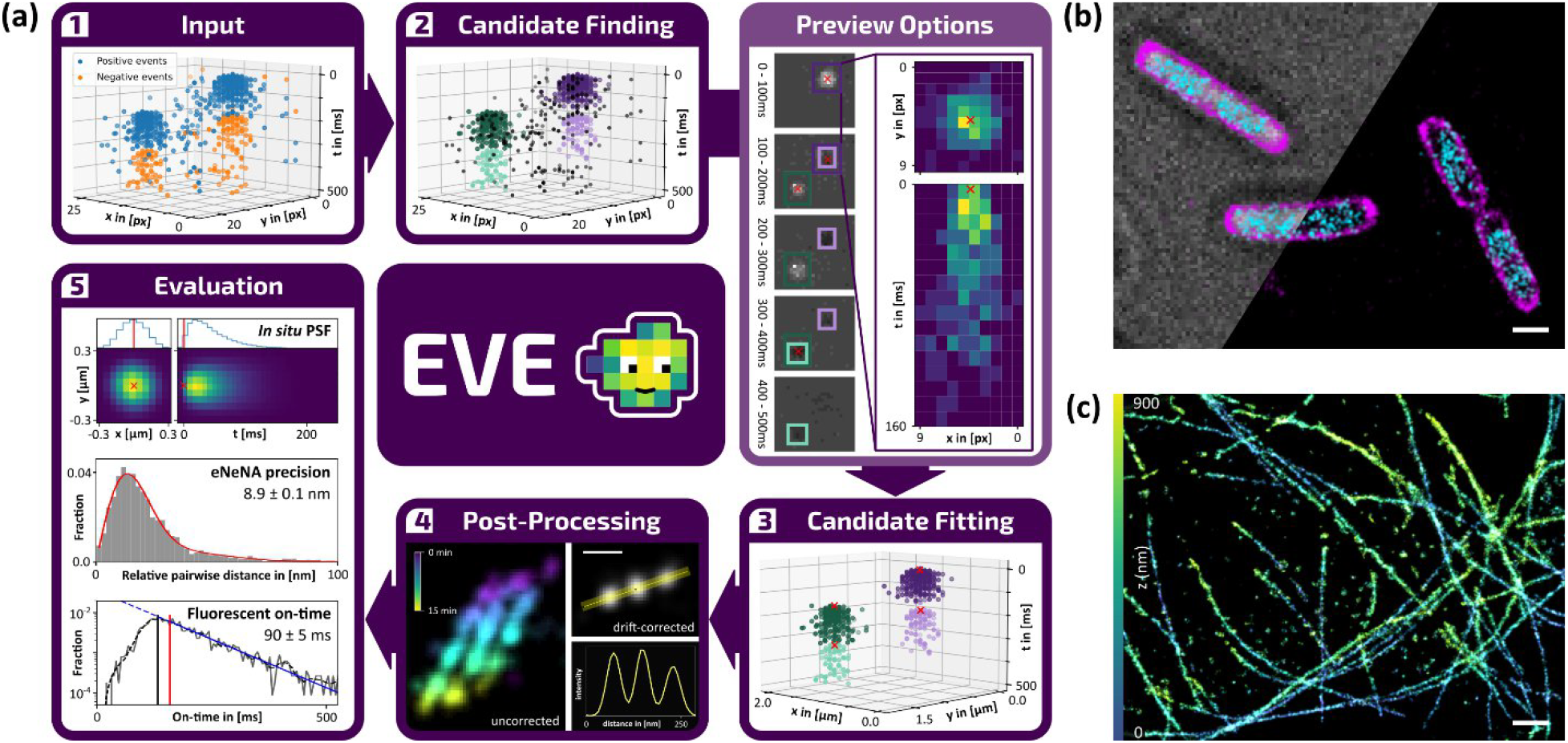
a) Working principle of EVE. First, the raw input, here an exemplary eveSMLM recording of a DNA-PAINT nanoruler (Supplementary Methods, Supplementary Note 2), is loaded and displayed as point-cloud data (1). The “Candidate finding” module (2) extracts candidate clusters belonging to single emitters; and the “Candidate fitting” module (3) determines exact x,y(,z)(,p),t coordinates for each candidate cluster. Various preview options can be used to assess the performance of both steps. The results can be post-processed (4), e.g. filtered, drift-corrected or visualized (scale bar 100 nm). Additionally, EVE offers multiple evaluation options (5), e.g. determining the *in situ* PSF, obtaining the ensemble localization precision, or estimating the fluorescent on-time distribution (Supplementary Note 1). (b) Multiplexed recording of *Eschericia coli* cells that endogenously incorporate mEos3.2-A69T-labeled RNA polymerase (cyan) and were imaged by photoactivated localization microscopy (PALM), combined with a read-out of their Nile Red-stained membrane (magenta) by point accumulation in nanoscale tomography (PAINT) imaging. Top-left: data overlaid on sCMOS-captured brightfield; bottom-right: isolated localization data. Data was processed by Eigenfeature-based (mEos3.2-A69T) and frame-based (Nile Red) candidate finding, respectively, and logarithmic Gaussian candidate fitting (Supplementary Methods, Note 1 and 2); scale bar 1 μm. (c) Astigmatism-based three-dimensional direct stochastic optical reconstruction microscopy (dSTORM) recording of the α-tubulin network of COS-7 cells visualized by AF647-labeled antibodies. Data was processed by Eigenfeature-based candidate finding and astigmatic-Gaussian candidate fitting (Supplementary Methods, Note 1 and 2), scale bar 1 μm, color represents 900 nm z-range of the recording.

The subsequent “Candidate Fitting” module (Figure 1a, box 3) takes the list of candidate clusters and determines the x,y(,z)(,p),t-localization for each candidate. The various fitting routines can be flexibly combined with all finding routines and either provide a combined spatiotemporal fit, or assess the spatial and temporal information separately (Supplementary Note 1). Next to analyzing the point-cloud data directly it is possible to process the data e.g. using only first events per pixel or the mean time delay between events (Supplementary Note 1).

The “Post-Processing” and “Evaluation” modules (Figure 1a, box 4, 5) offer several routines as exemplary shown for DNA-PAINT nanoruler data (Supplementary Methods). Next to technical tools such as filtering, drift-correcting or displaying the data, EVE can measure the fluorescent on-time of the emitters and the NeNA^6^ localization precision by a modified event-based Nearest Neighbour Analysis (eNeNA) (Supplementary Note 1). Remarkably, EVE’s analysis achieving noise-free candidate finding and temporally precise candidate fitting, combined with the point-cloud nature and static signal repression of eveSMLM, allows the straightforward retrieval of *in situ* PSFs, an involved process for camera-based SMLM^7^ (Extended Figure 1a).

Finally, EVE is an open software framework that can easily be extended: New routines can be added as python code of predefined structure (Supplementary Note 3). EVE automatically detects and displays them in its graphical user interface.

We demonstrate EVE with a multiplexed recording of the membrane and RNA polymerase of *Escherichia coli* cells; and with a three-dimensional recording^8^ of the α-Tubulin network in COS7-cells (Figure 1b, c, Supplementary Methods). Furthermore, we assess the performance of EVE and its routines. Finding routines were optimized during development, resulting in an excellent, linear scaling run-time (Extended Figure 1b). Among the fitting routines, the logarithmic Gaussian provides the best localization precision (Extended Figure 1c), an intuitive result considering that consecutive events per pixel are triggered by a logarithmically increasing threshold.

**Extended Figure 1:**
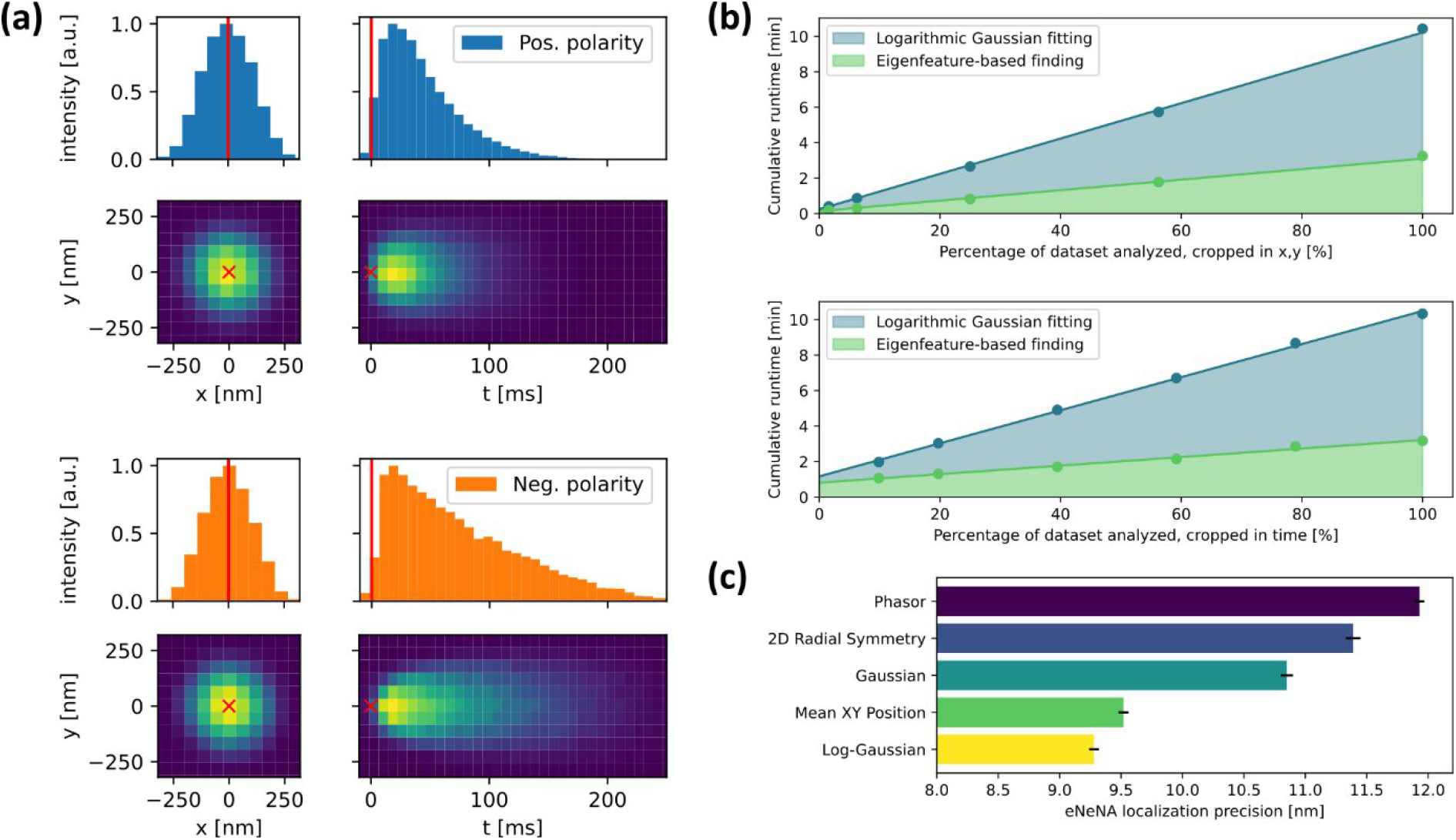
Assessing the performance of EVE using an exemplary DNA-PAINT nanoruler data-set (Supplementary Methods) (a) *In situ* PSF retrieval: By superimposing all candidates, the average x,y,t-PSF can be determined for both positive and negative event polarities, the blue and orange histograms display the distributions along the PSF center in y and t direction, respectively. The red cross or line mark the fitted coordinates used for superposition. (b) Run-time analysis of EVE: 100% of the data correspond to in total 173177 candidates found. The run-time shows a fast performance of EVE, completing the data analysis in the order of minutes and scaling linearly with the size of the data-set, measured by cropping the data in x,y (top) or cropping the data in time (bottom). (c) eNeNA localization precision of the different 2D fitting methods implemented in EVE, the logarithmic Gaussian outperforms all other methods.

In summary, EVE is a powerful, ready-to-use software package for eveSMLM data analysis, making the use of EBS a state-of-the-art method for SMLM imaging and easily accessible to any interested user. We envision EVE to serve as a versatile community platform due to its expandable, open-source infrastructure, facilitating further exploration of EBS use and helping to realize their full potential in SMLM imaging.

## Code availability

The software is available as a PIP package (https://pypi.org, eve-smlm) and can be accessed on github, https://github.com/Endesfelder-Lab/EVE-software. The scientific background of the implemented analysis methods, a user manual and the instructions for developers are distributed with the software and additionally provided as Supplementary Notes 1, 2 and 3.

## Data availability

All figure data is provided with the software on github, https://github.com/Endesfelder-Lab/EVE-software. Full source data of all recordings is available on Zenodo (doi: 10.5281/zenodo.13269600; https://zenodo.org/records/13269600).

## Acknowledgements

This work was financially supported by start-up funds at the University of Bonn (U.E.), an Alexander von Humboldt foundation fellowship (K.J.A.M.), by method development and open call funds of TRA Life and Health and the TRA Modeling (U.E.) and an Argelander Starter Kit (K.J.A.M.) at the University of Bonn as part of the Excellence Strategy of the federal and state governments, an Add-on Fellowship for Interdisciplinary Life Science by the Joachim Herz Stiftung (L.W.), a RISE travel stipend by DAAD/MITACS (K.H.) and a travel stipend from the Global Education Office at the University of New Mexico (J.G.). I.I, C.C, and M.A. acknowledge financial support under the programme ‘Investissements d’Avenir’ launched by the French Government; by the French Agence Nationale de la Recherche, project ABC4M under reference ANR-20-CE45-0023; and from Région Ile-de-France, DIM-ELICIT, OPI project.

We would also like to thank all members of the U.E. and I.I. laboratories for the valuable discussions.

## Author contributions

Conceptualization: K.J.A.M. and U.E. Data curation: L.W., K.J.A.M., C.C., M.A., F.V., Formal analysis: L.W., K.J.A.M., F.V., K.H., J.G., M.A., Funding acquisition: L.W., K.J.A.M., U.E., I.I. Investigation: L.W., K.J.A.M., C.C., M.A., F.V., K.H., J.G. I.I., and U.E, Methodology: L.W., K.J.A.M., C.C., F.V., K.H., J.G., I.I., and U.E. Project administration: K.J.A.M., U.E. Software: L.W., K.J.A.M., K.H., J.G., Supervision: K.J.A.M., I.I. and U.E. Visualization: L.W., K.J.A.M., K.H., and J.G. Writing—original draft: L.W., K.J.A.M., U.E. Writing—review and editing: all authors.

