## Supplementary Methods and Supplementary Note 1-3 for "EVE is an open modular data analysis software for event-based localization microscopy"

#### *E. coli* sample preparation

*E. coli* MG1655 rpoC:mEos3.2-A69T (CamR) was grown o/n from -70 °C glycerol stocks in M9 minimal medium (Merck) at 37 °C under constant agitation. This culture was subcultured 1:100 in EZRDM (EZ rich defined medium; Teknova) complemented with 1:100 20 % glucose, and grown for 24h, after which it was subcultured the same. This culture was grown for 3 hours, and the cells were directly fixed by adding 150 µL 37 % glycerol (pre-warmed to 37 °C) to 1350 µL culture and gentle agitation, and this fixation was continued for 20 minutes at 37 °C. The cells were washed twice with 500 µL PBS at 37 °C by centrifugation at 9000 g, and finally concentrated in 200 µL PBS. This sample was placed on an 8-chamber Ibidi slide, which was beforehand prepared as follows: the slide was cleaned by 15 minutes incubation with 1 M KOH, washed twice with distilled water, incubated with poly-L-lysine (PLL) for 20 minutes, washed twice with distilled water. After incubation of the sample on the slide, the sample was washed twice with PBS before imaging.

#### Imaging and Analysis of DNA-PAINT origami and *E. coli* cells

A custom laser-based fluorescence microscopy setup was used for imaging. 561 nm laser light (gem 561 1000, Laser Quantum) was modulated by an AOTF (G&H; AOMO 3080-125 & AODR 1080AF-AINA-1.0 HCR) and directed via a reflective collimator (RC04FC-P01, Thorlabs) into a custom fibre (70x70 µm multimode square core fibre, NA 0.22, FC/PC connectors, CeramOptec). 405 nm laser light (LBX-405-1200-HPE-PPA, RPMC) was co-aligned and also directed into the same fibre. The fibre output was collimated via a Multimode Collimator (F950FC-A 350-700nm, Thorlabs), and the beam was expanded via a set of lenses (LB1471-A-ML and LA4725-A, Thorlabs), and cleaned up via a ZET405/488/561/640 filter (QuadLineLaserClean-Up, AHF analysentechnik). This beam was then focused via an achromatic lens (AC254-400-A-ML, Thorlabs) and a dichroic mirror (ZT405/488/561rpc, Chroma, Bellows Falls, VT, USA) embedded in a commercial inverted microscope body (Nikon Eclipse Ti-E, Nikon, Tokyo, Japan) equipped with a focus stabilization system (Perfect focus system) on the back-focal plane of a 60x Apochromat TIRF 1.49 NA objective (Nikon). Emission from the sample passed via the objective and dichroic mirror through a filter (ZET405/488/561m-TRF, Chroma) and a 4f system via two convex achromatic lenses (AC508-100-A, Thorlabs), and optionally passed through an emission filter (ET610/75m, Chroma). The light was then directed either towards a Prime BSI sCMOS camera (Teledyne Photometrics, Tucson, AZ, USA; 107 nm pixel size), or towards an event-based sensor (Metavision Gen4.1-HD EVK, Prophesee; 80 nm pixel size). The microscope, camera, and peripherals were controlled via MicroManager 2.0, and laser triggering was controlled via a TriggerScope 4 (Advanced Research Consulting, Newcastle, CA, USA).

For *E. coli* rpoC imaging, the sample was illuminated with ~2 kW/cm2 561 nm laser, while the 405nm laser was increased manually from 0 to ~8 W/cm2 to have a low and steady photoactivation rate, until no new signal appeared. For the Nile Red imaging, the buffer was exchanged for PBS containing 12.5 nM Nile Red, and the sample was illuminated with ~2 kW/cm2 561 nm laser and the data was recorded for 3 minutes.

For *E. coli* Nile Red analysis, the frame-based finding method (Detection threshold = 3.0, Exclusion radius = 4.0, Min. Radius = 1.25, Max. Radius = 4.0, Frame time (ms) = 100.0 and Candidate radius = 4.0) was used with the same parameters for positive and negative event polarities separately. For *E. coli* rpoC analysis, the Eigenfeature-based finding method (*Positive polarity:* Linearity cutoff = 0.7, Maximum Eigenvalue cutoff = 45, Number of neighbours = 50, Ratio ms to px = 40, DBSCAN epsilon = 4 and DBSCAN nr. neighbours = 8; *Negative Polarity:* Linearity cutoff = 0.7, Maximum Eigenvalue cutoff = 75, Number of neighbours = 50, Ratio ms to px = 40, DBSCAN epsilon = 4 and DBSCAN nr. neighbours = 5) was used. Logarithmic Gaussian fitting (Time fit routine = Lognormal CDF (first events, weighted), distribution = 2D histogram of x,y positions, expected width = 150.0, fitting tolerance = 4.0) was used for both Nile Red and rpoC.

For DNA-PAINT nanoruler (80RG, Gattaquant) imaging, the sample was illuminated with ~2 kW/cm2 561 nm laser in TIRF mode by moving the focused spot to the side of the back-focal plane of the objective, and ~20 minutes of data was recorded.

For DNA-PAINT nanoruler analysis, the Eigenfeature-based finding routine (*Positive polarity:* Linearity cutoff = 0.7, Maximum Eigenvalue cutoff = 18, Number of neighbours = 50, Ratio ms to px = 10, DBSCAN epsilon = 3 and DBSCAN nr. neighbours = 12; *Negative Polarity:* Linearity cutoff = 0.7, Maximum Eigenvalue cutoff = 18, Number of neighbours = 20, Ratio ms to px = 18, DBSCAN epsilon = 3 and DBSCAN nr. neighbours = 8) was combined with logarithmic Gaussian fitting (Time fit routine = Lognormal CDF (first events, weighted), distribution = 2D histogram of x,y positions, expected width = 150.0, fitting tolerance = 1.0; same parameters for both event polarities).

### $\alpha$-tubulin sample preparation

African green monkey kidney cells (COS-7) were cultured at 37 °C and 5 % $CO_2$ in DMEM medium containing glutamax (Gibco No. 31966-047), 10 % fetal bovine serum (FBS, Gibco No. A3840401) and 50 U/ml penicillin and 50 µg/ml streptomycin (Gibco No. 15140-148). For experiments, cells were plated on 25 mm diameter glass coverslips (type 1.5, Marienfeld) placed in six-well plates containing culture medium with 2 % FBS at low density, and fixed on the following day in 0.1 M sodium phosphate buffer (PB), pH 7.4, containing 4 % paraformaldehyde (PFA), 0.2 % glutaraldehyde, 1 % sucrose, at 37 °C for 10 minutes, followed by three rinses in PBS. Cells were permeabilized with PBS containing 0.1 % Triton X-100 for 10 minutes and rinsed three times with PBS prior to immunolabeling. For the labeling, the cells were incubated for 1 hour at 37 °C with 1:300 mouse anti-alpha-tubulin antibody (Sigma Aldrich, T6199) in PBS + 1 % BSA. This was followed by three washing steps in PBS + 1 % BSA, incubation for 45 minutes at 37 °C with 1:300 goat anti-mouse AF647 antibody (Life Technologies, A21237) diluted in PBS + 1 % BSA and three more washes in PBS. Finally, the cells were post-fixed with 3.6 % formaldehyde for 15 min in PBS. The cells were washed in PBS three times and then reduced for 10 minutes with 50 mM $NH_4Cl$ (Sigma Aldrich, 254134), followed by three additional washes in PBS.

The fluorescent beads sample used for the calibration was prepared by diluting dark red 40 nm fluorescent beads (F10720, Thermo Fisher) with a dilution factor of 1:107 in phosphate buffered saline (PBS) and allowing them to deposit on a coverslip.

### Imaging and Analysis of $\alpha$-tubulin samples

A custom-built microscope with a RM21 body and a MANNZ nano-positioner (Mad City Labs) was used. The illumination and fluorescence collection were done with a Nikon 100x 1.49NA APO TIRF SR oil immersion objective. The excitation was performed via a 638 nm laser (LBX-638-180, 180 mW, Oxxius) and consisted of a vertical Gaussian beam with a diameter of 30 µm in the object plane. A full multiband filter set (LF405/488/561/635-A-000, Semrock) was used to separate and clean the illumination and the fluorescence. The fluorescence was sent in the detection module and recorded on the event-based sensor (EVK V2 Gen4.1, Prophesee). We used two afocal doublets (Thorlabs) to adjust the pixel size to 67 nm in the object plane. The 3D information was encoded by astigmatism using a cylindrical lens (focal length = 500 nm, Thorlabs) to achieve a separation of the x and y widths minima of 400 nm.

3D SMLM acquisitions on COS-7 cells were done using a continuous excitation with an irradiance of 5 kW/cm$^2$. We used a dSTORM buffer composed of 100 mg/ml glucose, 3.86 mg/ml MEA, 0.5 mg/ml glucose oxidase and 1.18 µl/ml catalase in PBS. After a pumping phase of a few minutes, the acquisitions were started, and stopped after 25 minutes. A low power continuous 405 nm excitation was also added during the second half of the acquisition to increase the density of detections.

For the calibration acquisitions, the excitation power was modulated with a square signal at a frequency of 30 Hz and with a duty cycle of 0.5. A series of acquisitions was performed with a 100 nm nanopositioner z position difference between each plane. These positions were converted to the corresponding positions of the focus plane by applying a focal shift correction coefficient of 0.75, as previously calibrated with a sample of known geometry [1].

All calibration files (each containing data aquired at different z-positions) were analyzed with EVE, treating all event polarities equally and performing a frame-based finding (Detection threshold = 3.0, Exclusion radius = 4.0, Min. Radius = 1.25, Max. Radius = 4.0, Frame time (ms) = 6000 and Candidate radius = 4.0) and 2D Logarithmic Gaussian fitting (Time fit routine = Average time, distribution = 2D histogram of x,y positions, expected width = 200.0, fitting tolerance = 10.0). The maximum size of a bounding box was adjusted to the frame time of 6000 ms in the advanced settings. For each calibration file analyzed, the x,y-widths of all localizations were then averaged and finally all average localizations were fitted with a fourth degree polynomial depending on their z-position. Hereby, the first 4 datapoints were excluded from the fit due to strong spherical aberrations.

The full dataset was then analyzed with Eigenfeature-based finding (*Positive polarity:* Linearity cutoff = 0.7, Maximum Eigenvalue cutoff = 23.5, Number of neighbours = 30, Ratio ms to px = 2.5, DBSCAN epsilon = 3 and DBSCAN nr. neighbours = 15; *Negative Polarity:* Linearity cutoff = 0.7, Maximum Eigenvalue cutoff = 17.2, Number of neighbours = 30, Ratio ms to px = 6.0, DBSCAN epsilon = 3 and DBSCAN nr. neighbours = 15) and astigmatic logarithmic Gaussian fitting (Time fit routine = Average time, distribution = 2D histogram of x,y positions, expected width = 200., fitting tolerance = 5.0; same parameters for both event polarities).

### Hardware and software environment

All analysis, including the performance tests, was performed on a 64-bit PC equipped with a i5-12400 12th gen Intel Core processor (6 cores, 12 threads) at 2.50 GHz, 32 GB RAM (3600 MT/s), NVIDIA RTX 4070 Ti (7680 cores, 12 GB memory), on a PRIME B660M-K D4 motherboard. EVE was run via Python 3.9.18 running in a Conda environment in Windows 11 Pro, and analysis was performed with processing parallelization turned on.

### Supplementary Note 1: Analysis methods implemented in EVE

#### Contents


- Introduction
- Finding
  - Eigenfeature-based finding
  - DBSCAN-based cluster finding
  - Frame-based finding
- Fitting
  - Fitting distributions
  - Mean X,Y position
  - 2D Gaussian
  - 2D Logarithmic Gaussian
  - 3D Astigmatic Gaussian
    - Model function
    - Calibration
    - Estimating the axial position
  - Phasor-based Fitting
  - Radial Symmetry Fitting
  - Temporal fitting
    - Lognormal CDF fitting
    - Temporal Gaussian fitting
- Post-processing and Evaluation
  - Polarity matching post-processing
    - Polarity matching
    - Localization precision
    - Estimation of the emitter fluorescent On-time
  - Drift correction
  - Visualisation

### Introduction

EVE is a software platform developed for the analysis of single-molecule imaging data captured by event-based sensors. The software is methodically divided into three integral modules, each serving a distinct purpose in the data analysis pipeline for event-based single-molecule data.

1. **Candidate Finding:** This initial module is responsible for identifying and isolating potential single-molecule signals within the event data. It effectively filters the raw input to extract candidate events for further analysis.
2. **Candidate Fitting:** Once candidates are identified, this module precisely (sub-pixel) localizes the single molecules spatiotemporally.
3. **Post-Processing and Evaluation:** This module includes various routines to modify and interpret the data.

### Finding

#### 1. Eigenfeature-based finding

The Eigenfeature analysis finding methodology is based on spectral clustering methods, which are used in 3D point-cloud data classification such as LiDAR [2], [3], [4], [5]. Spectral clustering conceptually works as follows: First, for each point in a point cloud, a subset of the full point cloud is extracted, either by looking in a certain (multi-dimensional) radius, or alternatively by extracting the nearest N neighbors. Secondly, the corresponding covariance matrix of this subset is calculated and attributed to the original point. Finally, from this covariance matrix, the Eigenvalues are extracted. For event-based single-molecule data, three Eigenvalues are extracted, corresponding to the three dimensions (two spatial, one temporal). This is repeated for all points in the point cloud.

The extracted Eigenvalues contain information about local geometric features around each event. These geometric features are called 'Eigenfeatures'. In EVE, we use Eigenfeatures to discriminate points belonging to single-molecule emissions from noise. In EVE, Eigenvalues are determined by assessing the nearest N neighbors of each event in the event point cloud (i.e. the complete dataset). N should be chosen to be roughly equal to the expected size of a single-molecule emission event, in combination with a temporal-to-spatial factor so that temporal and spatial distances can directly be compared. We then assess two Eigenfeatures:

1. The first Eigenfeature is the maximum Eigenvalue for each event in the point cloud. The maximum Eigenvalue is high if no local cluster features are present, and low if a local cluster feature is present (see Supplementary Figure 1). Therefore, only events with an associated low maximum Eigenvalue can be considered to be part of single-molecule localizations. The cut-off Eigenvalue for a dataset can be determined by finding valleys via a wavelet transformation (Python scipy package).
2. The second Eigenfeature is the 'Dimensionality' [2], [3]. The dimensionality can be derived by comparing the magnitude of the three extracted Eigenvalues of each event; if they are similar (i.e. $\lambda_1 \approx \lambda_3 \approx \lambda_3$), the local feature represents a volumetric structure, whereas if one Eigenvalue is much larger than the other (i.e. $\lambda_1$»$\lambda_2$,$\lambda_3$), the local feature has a one-dimensional shape. In EVE, we use this to discriminate between single-molecule clusters (which are three-dimensional in shape when assessing time as the third dimension) and artifacts created by hot pixels (which are one-dimensional in the time dimension).

These two Eigenfeatures are combined to discriminate between signal and noise in EVE: if the first Eigenfeature is a lower value than the cutoff, and if the second Eigenfeature is considered volumetric, the single event is considered signal, otherwise noise. Downstream of this discrimination, a DBSCAN clustering routine is performed to further filter out spurious mis-labeled events. The events contained in these clusters are the candidates passed on for sub-pixel localization (Fitting).

#### 2. DBSCAN-based cluster finding

Density-based spatial clustering of applications with noise (DBSCAN) is an algorithm which clusters points based on their local density, and is routinely used in SMLM data analysis [6]. To save computation time for the DBSCAN algorithm, we remove most noise in two steps. First, (only in the case of analysing positive and

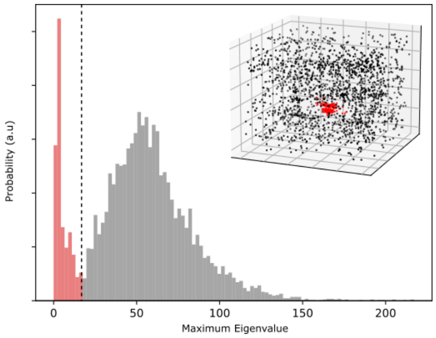


**Supplementary figure** 1: Typical histogram depicting the maximum Eigenvalue of each event of eveSMLM data which is used to discern single-molecule emissions from noise as noise typically has higher values. With a threshold of a maximum Eigenvalue lower than 17 (dashed line), the single-molecule data can be separated (in red highlighted events in the insert).

negative events together), we filter out events that show either an on-to-off-event or off-to-on-event sequence per pixel (Supplementary Figure 2 a, b). This is based on the rationale that single-molecule emissions usually trigger a sequence of multiple, consecutive on-events while turning on (and a sequence of multiple off-events when going dark) and thus trigger several thresholds and therefore show event sequences of the same polarity. Noise, on the other hand, is more likely to fluctuate around an on average constant background level, resulting in polarity-switching on-to-off and off-to-on event sequences. Second, for each event, a neighbor count is performed within a set spatiotemporal radius and compared against the average data density of the entire dataset (Supplementary Figure 2 c). Only events which have at least N (user-definable) times more neighbors than average remain, others are discarded. This 'high-density dataset' is then processed via DBSCAN with user-definable radius and minimum cluster points, to obtain single clusters for each single-molecule emission (Supplementary Figure 2 d). The bounding boxes of this cluster (optionally plus an additional area of user-defined spatiotemporal size) are then used to extract all, unfiltered events (i.e. raw events except hot-pixels, which are removed by a filter for consecutive $> 30$ positive or negative events) in this bounding box. These final candidate are passed on for sub-pixel localization.

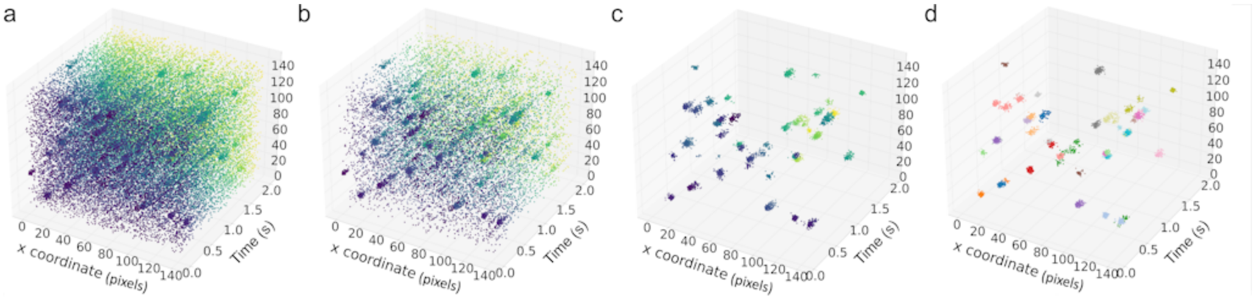


**Supplementary figure** 2: Schematic representation of the DBSCAN-based finding routine. a. Raw event data in a certain spatiotemporal area. b. Consecutive event-filtering, where only on-to-on and off-to-off sequences of events remain. c. Filtering for events that have a density higher than 1.5 times the average density in a [240 nm; 40 ms] radius. d. Clusters obtained when running the DBSCAN algorithm on the data in c with radius of 6 and minimum cluster points of 15.

### 3. Frame-based finding

The frame-based finding methodology is an adapted version of the frame-based analysis used in [7]. It naively reduces event-data to images using 'frames' of e.g. 100 ms. In these frames, standard image blob detection methods can be employed – in our implementation, an image wavelet segmentation method is used

[7], [8]. This segmentation method reports a single spatial location (in pixel-units), and a user-defined area is extracted around this location. All events in this spatial area, in the temporal region defined by the frame, are passed on for sub-pixel localization.

### Fitting

All fitting methods aim to determine the x-, y- (,z-) and t-coordinates for each candidate cluster. Since a candidate cluster consists of a three-dimensional point cloud of events (see Supplementary Figure 3 a), conventional single-molecule localization methods that process image data cannot be used directly. Various event-based fitting methods are implemented in EVE, which can be divided into spatial, temporal and spatio-temporal fitting method types. The first two method types can be flexibly combined to obtain the final localization, whereas the third method type directly estimates both temporal and spatial coordinates together.

#### 1. Fitting distributions

Besides directly analyzing the event point cloud data associated with a candidate (Supplementary Figure 3 a), it is also possible to reduce the data to a two-dimensional distribution, e.g. by counting all events per pixel (Supplementary Figure 3 b), by only taking the time of the first events per pixel (Supplementary Figure 3 c) or by calculating the mean time delay between all events per pixel (see Supplementary Figure 3 d). These distributions can then be fitted by a conventional two-dimensional function, e.g. a Gaussian, to obtain the x,y(,z) localization.

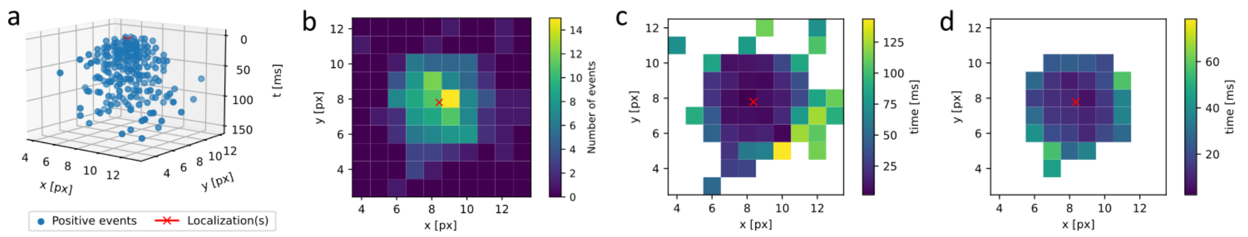


**Supplementary figure** 3: Fitting distributions that can be derived from the event data. a. Event point cloud belonging to a candidate. b. Total number of events per pixel. c. Time of the first event per pixel (pixels with 0 events cannot be evaluated). d. Mean time delay between all events per pixel (pixels with less than 2 events cannot be evaluated).

Within EVE, we separated the creation of the two-dimensional distribution from the fitting of this distribution. This allows a very flexible combination of both, method and distribution, to achieve the most accurate localization result by fully evaluating and exploiting the information stored in a candidate cluster. Two-dimensional fitting routines can be chosen from: Mean X,Y position; Gaussian; logarithmic Gaussian; astigmatic Gaussian and Phasor routines. Two-dimensional distributions can be chosen from: The time of the first event; the total number of events; the average/median time of all events for each pixel; the average/minimum/maximum time delay between events for each pixel.

### 2. Mean X,Y position

The average X,Y fitting method calculates the average x,y position for the chosen two-dimensional distribution, the uncertainties of the spatial x,y estimators are determined by the standard deviation in x and y. A temporal fitting method has to be selected separately.

### 3. 2D Gaussian

Analogous to conventional frame-based SMLM [9], [10] a 2D Gaussian least-square fit with variable x- and y-width ($\sigma_x$, $\sigma_y$) is implemented in EVE:

$$pdf(x,y) = A_1 \cdot e^{-\frac{(x-x_0)^2}{2\sigma_x^2} + \frac{(y-y_0)^2}{2\sigma_y^2}} + A_2$$

$A_{1-3}$ are the amplitudes of each term. The ($x_0$, $y_0$) - localization uncertainties given by the fit are used as a tolerance measure to discard imprecise fits where the fit uncertainty is larger than a user-definable factor times the pixel-size. A temporal fitting method has to be selected separately.

### 4. 2D Logarithmic Gaussian

Due to the logarithmic nature of consecutive events a 2D logarithmic Gaussian fit was implemented similar to the 2D Gaussian fit:

$$pdf(x,y) = A_1 \cdot log(1 + e^{-\frac{(x-x_0)^2}{2\sigma_x^2} + \frac{(y-y_0)^2}{2\sigma_y^2}}) + A_2$$

Again, the ($x_0$, $y_0$) - localization uncertainties given by the fit are used as a tolerance measure to discard imprecise fits where the fit uncertainty is larger than a user-definable factor times the pixel-size. A temporal fitting method has to be selected separately.

### 5. 3D Astigmatic Gaussian

This method is used to adjust 3D eveSMLM data generated by inserting a cylindrical length into the detection path to introduce a slight astigmatism into the image [11]: While the PSF appears round and symmetrical in the focal plane, it is elongated in the x and y directions above and below the focal plane, respectively. In this method, spatial and temporal fitting are separated again, so that an additional temporal fitting method must be selected.

#### 5.1. Model function

A rotated elliptical Gaussian function with rotation angle $\phi$ is used to represent the astigmatic PSF:

$$pdf(x,y) = A_1 \cdot e^{-\frac{\hat{x}^2}{2\sigma_x^2} + \frac{\hat{y}^2}{2\sigma_y^2}} + A_2$$

where $A_{1,2}$ are amplitude terms, $\sigma_{x,y}$ are the imaged widths of the molecule along two perpendicular axes rotated by $\phi$ with respect to the x,y-axes and x,y are defined by:

$$\hat{x} = (x - x_0)cos(\phi) - (y - y_0)sin(\phi)$$

$$\hat{y} = (x - x_0)sin(\phi) + (y - y_0)cos(\phi)$$

Here, $x_0$ and $y_0$ again represent the sub-pixel coordinates of the molecule.

##### 5.2. Calibration

For calibration a sample containing immobile fluorescent beads is imaged at different z-planes. For each z-plane the average x- and y- width $\sigma_{\hat{x},\hat{y}}$ are determined and then fit by a fourth order polynomial:

$$\sigma_{\hat{x},\hat{y}}(z) = a_{\hat{x},\hat{y}} \cdot (z - c_{\hat{x},\hat{y}})^2 + d_{\hat{x},\hat{y}} \cdot (z - c_{\hat{x},\hat{y}})^3 + e_{\hat{x},\hat{y}} \cdot (z - c)^4 + b_{\hat{x},\hat{y}}$$

##### 5.3. Estimating the axial position

The axial position is estimated by minimizing the distance between the ratio of measured x- and y-width $\Sigma_{\hat{x}}/\Sigma_{\hat{y}}$ to the calibrated ratio curve $\sigma_{\hat{x}}(z)/\sigma_{\hat{y}}(z)$, thus by

$$z = \min_z \left( \left( \frac{\Sigma_{\hat{x}}}{\Sigma_{\hat{y}}} - \frac{\sigma_{\hat{x}}(z)}{\sigma_{\hat{y}}(z)} \right)^2 \right)$$

The z-uncertainty is determined via the inverse Hessian matrix returned by the minimization routine.

#### 6. Phasor-based Fitting

The phasor-based fitting routine is directly adapted from the pSMLM fitting for camera-based SMLM data [12]. Briefly, the algorithm converts the 2D distribution to two phase vectors (or phasors) by calculating the first Fourier coefficients in x and y. The angles of these phasors are used to localize the center of the event distribution.

This concept can be expanded to the third dimension if a temporal binning is employed as well as a spatial binning.

#### 7. Radial Symmetry Fitting

This fitting routine is an adapted version of the calculation of the radial symmetry centers for localization described in [13]. The general idea is to estimate the center of radially symmetric intensity distributions, by tracing lines parallel to the image gradients in each point, where the distance of all lines is minimal at the center. This approach does not require knowledge of the exact shape of the distribution. In the two-dimensional version, an array of event numbers per pixel is used instead of an array of pixel intensities. As shown in Supplementary Figure 3, the two-dimensional event distribution also shows a clear radial symmetry. The concept is then expanded to a three-dimensional version in which the events are translated into a three-dimensional histogram of event numbers per voxel. Again the point of maximum radial symmetry can be calculated via the image gradient.

#### 8. Temporal fitting

The aim of all temporal fitting methods is to get an accurate estimate for the true time of the brightness change. As the response time of sensor depends on imaging, sensor settings and pixel history and thus may be varying during the experiment, the time point of the emerging signal of a cluster as recorded by the sensor is taken as most consistent estimate of the true time.

#### 8.1. Lognormal CDF fitting

In this temporal fitting method, the cumulative number of events is fitted by a lognormal CDF to obtain the starting time $t_0$ of the candidate cluster. Throughout the data, we observe a wide variety of different temporal profiles for candidate clusters. We therefore chose the cumulative distribution function (CDF) of the lognormal function for fitting, as it allows a large freedom in shape, while a starting time $t_0$ can be estimated consistently.

$$\mathrm{CDF}(t) = \frac{A_1}{2} \cdot \left(1 + erf\left(\frac{\ln(t - t_s) - \mu}{\sqrt{2} \cdot \sigma}\right)\right) + A_2 \cdot (t - t_s) + A_3$$

To account for background noise, the fit consists of a first term containing the lognormal, and two additional terms that describe a linear dependency following from the assumption that the noise event rate is constant. $A_{1-3}$ are the amplitudes of each term, $erf$ is the Gaussian error function, $t_s$ is the shift of the fitting function, $\mu$ and $\sigma^2$ describe the mean and variance of the underlying normal function. The starting time of the cluster can now be estimated through the intersection of the maximum slope of the fit (gray curve in Supplementary Figure 4) and the background model (dashed line in Supplementary Figure 4).

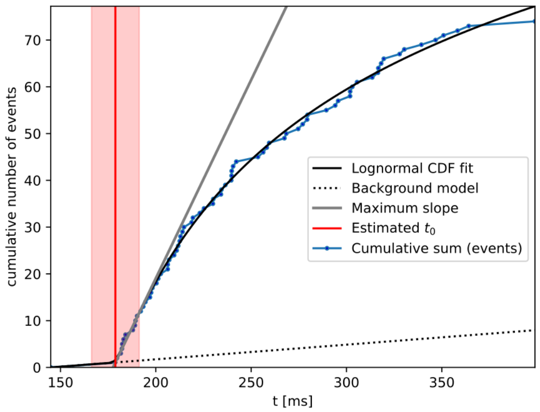


**Supplementary figure** 4: Exemplary representation of lognormal CDF fitting for time estimation. The cumulative number of events (blue) per cluster is fitted by a lognormal cumulative distribution function including a linear background correction model (black). The time (red) can be estimated via intersection of the maximum slope (gray) and background curve (dashed black line).

Instead of fitting all events, the fit can also be performed for the first events per pixel and candidate (green curve in the Supplementary Figure 5), with both fits yielding similar time estimates. However, it should be noted that the first events are very sensitive to the accuracy of the candidate finding method and only contain precise timing information if they are not corrupted by noise signals that occur before the single-molecule emission (leading to the noise events being plotted as first event per pixel). To reduce the noise dependence, each first event can be weighted by the total number of events in each pixel per candidate cluster.

#### 8.2. Temporal Gaussian fitting

This fitting routine makes use of the dependency between the absolute change in brightness and the response speed of the sensor: Stronger changes in brightness trigger events earlier than lower changes in brightness, resulting in events with earlier timestamps near the center of the PSF, while events at the edges are triggered later. This radial symmetry of the time of the first events per cluster can be seen exemplified in Supplementary Figure 6 a. To determine the starting point of the cluster, the two-dimensional distribution of the first events is fitted by a two-dimensional Gaussian curve with variable x- and y-width as depicted

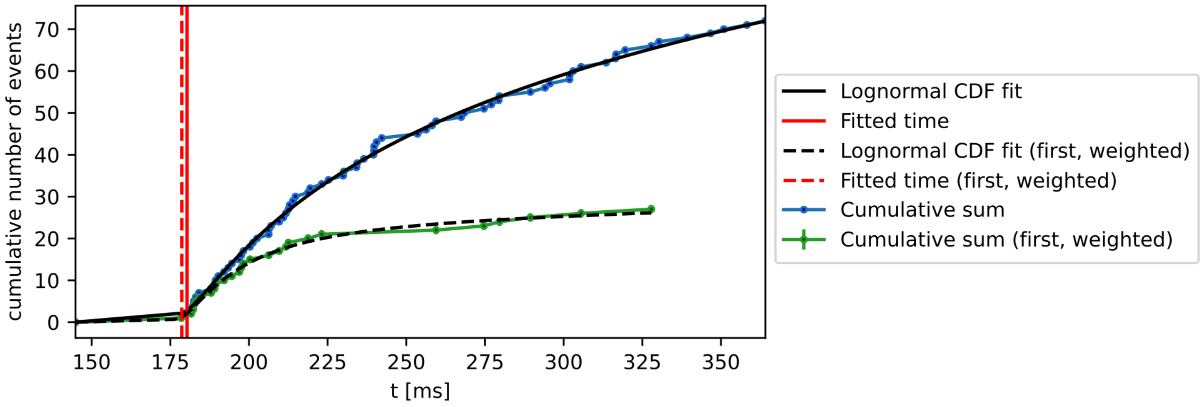


**Supplementary figure** 5: Exemplary representation of lognormal CDF fitting for time estimation of an off-switching event. Both the cumulative number of all events (blue) and the cumulative number of only the first events (green) can be used to estimate the time (red).

in Supplementary Figure 6 b. The starting time can now be estimated through the temporal peak of the Gaussian fit, which is reached in the center.

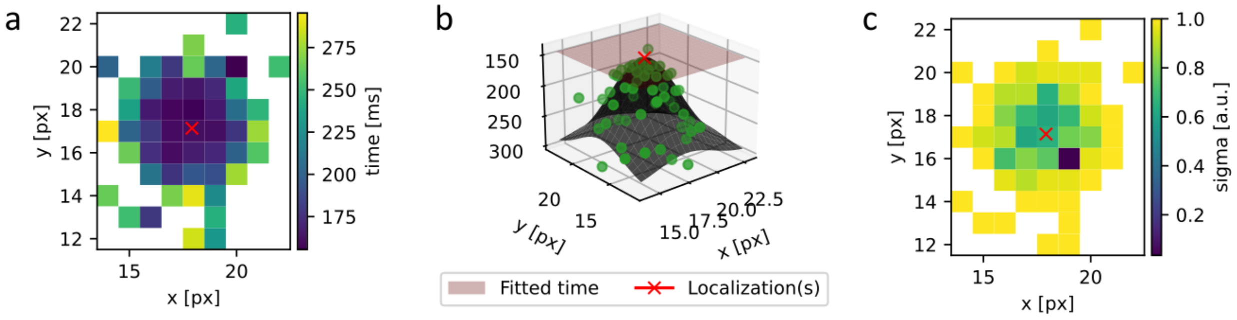


**Supplementary figure** 6: Exemplary representation of 2D Gaussian fitting for time estimation. a. Time of first event per pixel and candidate, central pixels are activated earlier. b. Two dimensional Gaussian fit (black) of all first events per pixel (green). c. Error map used as weight in the fit.

As noted earlier, the first events per candidate cluster are easily corrupted by noise. Therefore, it is only advised to use this fitting method, if the candidate finding method is stable against noise. To reduce the noise dependence, each first event is again weighted by the total number of events in each pixel per candidate cluster (see error map in Supplementary Figure 6 c).

### Post-processing and Evaluation

#### 1. Polarity matching post-processing

Due to the nature of event-based sensors to report changes in intensity rather than intensity itself, it is expected that a blinking single-molecule emitter will have both a “positive PSF” when turning on and a “negative PSF” when turning off or bleaching (i.e. rising edge and falling edge in intensity-profile). Thus, an eveSMLM dataset can be post-processed on this polarity information and double recording of each blinking emitter, and information can be extracted from this.

##### 1.1. Polarity matching

Matching of a “positive PSF” with its corresponding “negative PSF” searches for the closest negative PSF neighbor of the positive PSF. Bounds (default 200 nm radius, 0 to 1000 ms time) can be set by the user. Each positive PSF can only ever be linked to a single negative PSF and vice versa.

##### 1.2. Localization precision

In a static eveSMLM dataset (i.e. considering immobile molecules and ignoring sample drift), the underlying single-molecule location of the positive and negative PSF are identical, or $\Delta_{spatial} = 0$. However, in practice, the found localization difference between the positive and negative PSF ($\Delta_{spatial}$) is degraded by the combined localization precision of both positive and negative fitting routines. Nearest-neighbour analysis (NeNA) [14] is used to determine the average localization precision of all polarity-matched positive and negative PSFs in the dataset. For this, the $pdf(\Delta_{spatial})$ of all polarity-matched PSFs is fitted with the following function:

$$\mathrm{pdf}(\Delta_{spatial}) = A_1 \cdot \left( \frac{\Delta_{spatial}}{2 \cdot \sigma_{\mathrm{SMLM}}^2} \cdot e^{-\frac{\Delta_{spatial}^2}{4 \cdot \sigma_{\mathrm{SMLM}}^2}} \right) + A_2 \cdot \left( \frac{1}{\sqrt{2 \cdot \pi \cdot \omega^2}} \cdot e^{-\frac{(\Delta_{spatial} - d_c)^2}{2 \cdot \omega^2}} \right) + A_3 \cdot \Delta_{spatial}$$

The first term is the Rayleigh distribution from which $\sigma_{SMLM}$, the (mean) localization precision can be determined, the other two terms represent a Gaussian and a line noise correction, with $\omega$ being the Gaussian standard deviation characterizing the short-range correction term centered at $d_c$, and $A_{1-3}$ are the amplitudes of each term [14].

##### 1.3. Estimation of the emitter fluorescent On-time

In addition to spatial information, the polarity matching also provides temporal information about the emission cycle duration (i.e. *on-time*) of individual fluorophores. The *on-time* belonging to an entire dataset can be estimated.

The obtained duration between the positive-event PSF and negative-event PSF via polarity matching ($\Delta_{temporal}$) is broadly categorized in two regimes: 1. Photophysical *on-time* of a single fluorophore (i.e. caused by fluorophore bleaching (STORM, PALM) or target dissociation (PAINT)). This *on-time* is degraded by: 2. Sensor limitations: typical evePSFs have a temporal duration in the order of tens to hundreds of ms, even while the PSF signal is instantaneous. If the photophysical *on-time* of the single emitter is shorter than the evePSF formation, the positive PSF does not have time to fully form, and will as such not be localized correctly. Thus, low-value $\Delta_{temporal}$ are effectively removed from polarity-matching analysis.

The lifetime is estimated as follows (Supplementary Figure 7): the peak of the $\Delta_{temporal}$ pdf is determined via smoothing of the raw data with a Savitzky-Golay filter, after which an 'offset' is determined (default 20% higher than the temporal peak value). The raw data at times longer than this offset are fit with a combination of 1-3 exponential decays (user-definable).

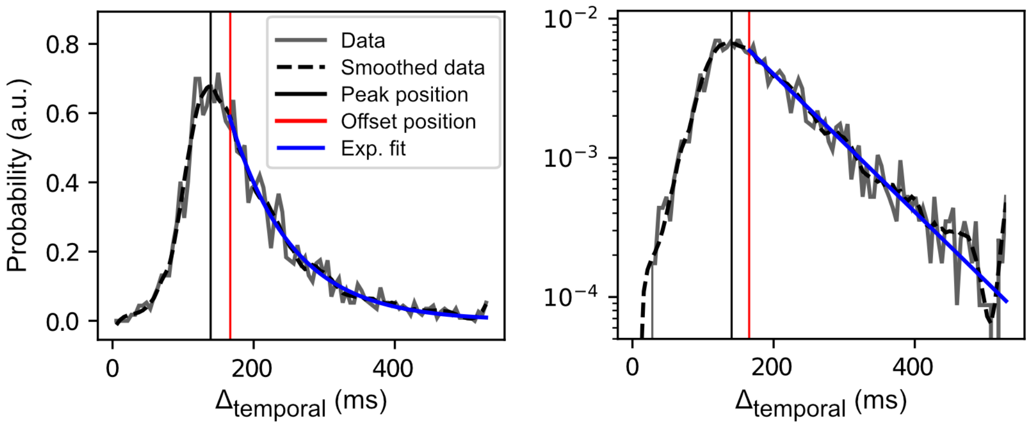


**Supplementary figure 7**: Showcase of the lifetime obtained via EVE from all emitter *on-times* of a DNA PAINT sample. A single exponential decay (blue line) is fitted to $\Delta_{temporal}$ pdf data (gray) on a linear (left) and logarithmic (right) y-axis. The exponential decay is fitted only in the regime where $\Delta_{temporal}$ is larger than the offset position (red). The single exponential decay fit has a half-time of 88 ± 3.5 ms.

### 2. Drift correction

Drift correction on the final localizations can be performed either via redundant cross correlation (RCC) [6] or entropy minimization (DME) [15]. Since RCC and DME are based on the concept of frames, a pseudo-frame-time should be provided for RCC/DME to run.

### 3. Visualisation

Visualisation of the final localizations [6] can be performed via individually rendered Gaussians with a global sigma, or with the sigma provided by the localization precision of each localization. Additionally, linearly interpolated 2D histograms can be created. For all methods, a visualisation pixel size should be provided.

### Supplementary Note 2: Software manual

#### EVE - General-purpose software for eveSMLM localization

#### About EVE

EVE is a graphical user-interfaced software package that provides several methods to localize emitters from single molecule localization microscopy (SMLM) experiments performed on event-based sensors (eveSMLM).

Event-based data differs fundamentally from conventional camera images. Unlike traditional sensors, event-based sensors only capture intensity changes, registering them as either positive (when the intensity surpasses a predefined threshold) or negative events (when the intensity drops below a predefined threshold). As a result, only a list of x and y pixel coordinate pairs is stored together with the detected event polarities (p) and timestamps (t).

EVE is designed to quickly and directly process and analyse event-based single molecule data. The event-based data analysis is divided into three main parts:
1. **Candidate Finding:** The complete event-list is searched for characteristic event clusters that are generated by blinking fluorophores. Potential candidate clusters are then extracted and returned for further processing.
2. **Candidate Fitting:** The x,y,(z),p,t-localization is determined for each candidate cluster.
3. **Postprocessing and Evaluation:** Various analytical routines to process and interpret the data.

EVE allows flexible combinations of different finding and fitting routines to optimize the localization results for the specific event-based dataset. Besides a variety of different finding and fitting algorithms, EVE also offers various preview options, visualisation tools and post-processing and evaluation functions. A detailed description of the algorithms can be found in **Supplementary Note 1: Analysis methods implemented in EVE**.

EVE is written in Python and structured in such a way, that is easy to implement and add new functionalties in form of new finding, fitting routines, etc. Details about this can be found in the **Supple-**

**mentary Note 3: Developer Instructions**. EVE can also be run via the command line interface, see `EVE_CommandLine.ipynb` for detailed information.

### Contents


- How to install and run EVE
  - Installation instructions
    - Optional
  - Running instructions
- Quick Start Guide
  - Set up the Processing Tab
  - Perform a Preview Run
  - Explore the Candidate Preview
  - Notes of best finding/fitting algorithms and parameters
  - Execute a full Run
  - Visualize the Localization Results
  - Apply a Drift Correction
    - Under Windows
    - Under Linux
  - The final Localization List
  - Estimate the Localization Precision
  - Metadata, run and result files
  - Command-line interface
  - Exemplary data


#### How to install and run EVE

This software was tested under Windows 10/11, Linux (Ubuntu 20.04) and macOS 14.5 (Sonoma). Besides the drift correction modules by entropy minimization (`Drift correction by entropy minimization [2D]` and `Drift correction by entropy minimization [3D]`) and the Gaussian visualization methods (`Gaussian_blurred with fixed sigma`, `Gaussian_blurred with locPrecision sigma`) that use pre-compiled dll-files and are therefore only running on Windows, everthing is running under MacOS, Linux and Windows. The software requires Python 3.9.

##### Installation instructions

Create a virtual environment using python 3.9, and install EVE with pip:

**With conda**
Install conda from the website (www.conda.io)
Run the following lines in the terminal:

```
conda env create --name EVE python=3.9
conda activate EVE
pip install eve-SMLM
```

**With virtualenv**

First, install python 3.9 from python.org.

Replace `PYTHON_PATH` by your python path, e.g. `/usr/bin` and `ENVIRONMENT_PATH` by the path to the virtual environments on your machine and follow the instructions below:

```
virtualenv -p PYTHON_PATH/python3.9 ENVIRONMENT_PATH/Eve
pip install eve-SMLM
```

##### Optional

EVE can read and process event-based data in `.npy` and `.hdf5` format. Additionally the `.raw` format of Prophesee can be used. If you have `.raw` data that you want to analyze you need to install the Metavision SDK from Prophesee beforehand (https://docs.prophesee.ai/stable/installation/index.html). We recommend using `.hdf5` whenever possible, as this is a hierarchical data format optimized for efficient saving and reading of large files.

#### Running instructions

Start the software by running `eve-SMLM` in the terminal. You can also download or clone EVE directly from github and install the required packages using the `requirements.txt` file. If you do this, you can start the software by running "GUI.py".

### Quick Start Guide

#### 1. Set up the Processing Tab

Running `eve-SMLM` or `GUI.py` in the terminal will open EVE's graphical user interface which you can see on the right side of the figure.

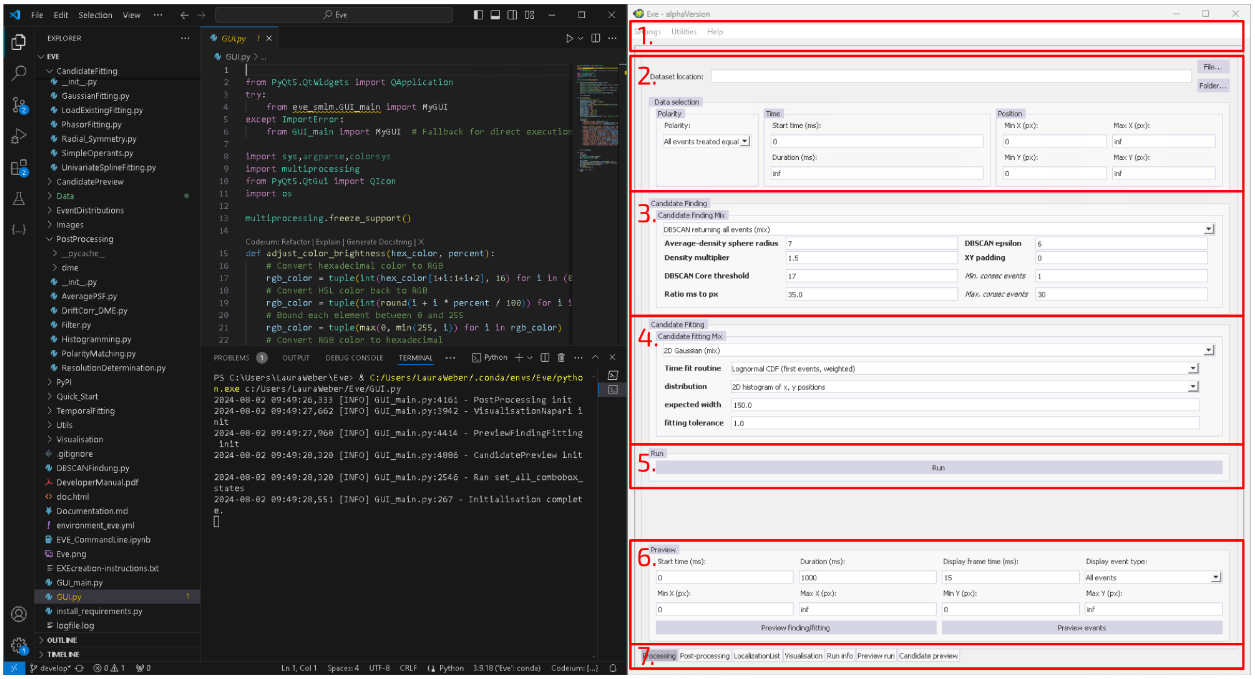


The main window has 7 major parts that are marked with red boxes and are described in more detail in the following.

1. **Menu bar:** By clicking on `Settings`, you can open the `Advanced settings`, save or load the current GUI configuration. Under `Utilities` you will find some additional functionalities to pre-process the raw event data files before processing them with Eve. Under `Help` you have access to all important information around EVE, such as the Users manual, the Developers manual and the Scientific information.

The folder Data\Nanoruler contains the GUI configuration (GUI_settings_Nanoruler.json) that we will use throughout this users manual. You can load the GUI configuration via Settings -> Load specific GUI contents and selecting the correct path. Now, open the Advanced settings, where you can adapt more advanced settings.

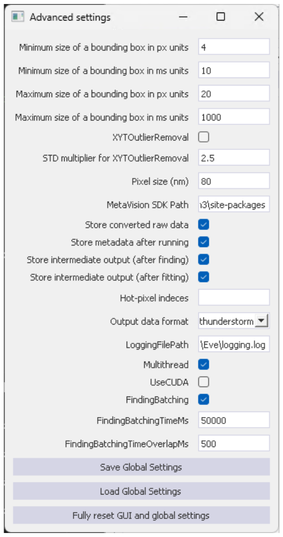


2. **Data to analyse:** Here, you specify the data that will be analysed in the following. You can either select a single file (in .npy, .hdf5 or .raw format) or a folder. If you select a folder all files in the folder will be analysed one after another.

The folder Data\Nanoruler contains an exemplary event-based acquisition of a DNA nanoruler (Nanoruler_Small.hdf5) which we will use in this tutorial. Fill the path entry field Dataset location: with the corresponding path to the nanoruler dataset.

In the Data selection box, you can now further specify which parts of the data should be analysed and how. You have different options for Polarity, Time and Position. Choose Pos and neg seperately as Polarity option while leaving the remaining settings unchanged. Thereby, you simply load the all events without temporal or spatial constraints. By selecting Pos and neg seperately all subsequent analysis steps will be run on the positive and negative events distinctly. If you change the Polarity option, the GUI will automatically adapt to your selection and show separate or combined options finding and fitting routines for positive and negative events seperatly.

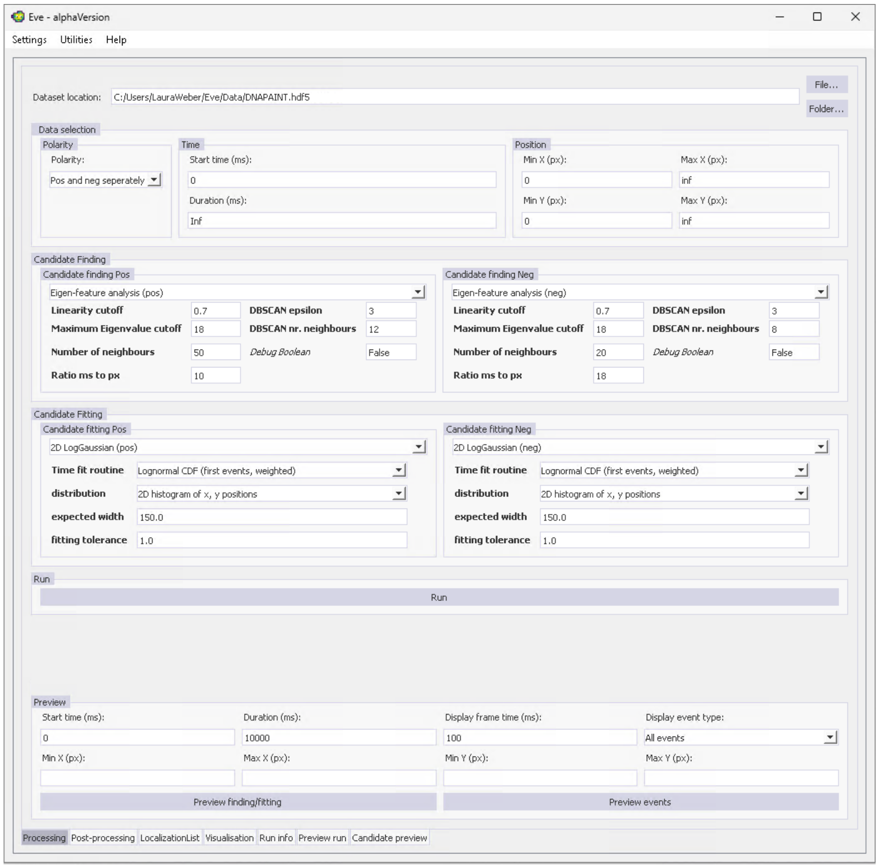


3. **Candidate Finding routine:** Here, you can select among different candidate finding routines. We will use `Eigen-feature analysis`, both for positive and negative events throughout this manual.
4. **Candidate Fitting routine:** Here, you can specify which fitting routines you want to use to get localizations for each candidate, cluster. We will use `2D Logarithmic Gaussian` throughout this manual, again for both polarities.
Everything is now ready for the first run. Before you start the first run, save the GUI settings (`Settings -> Save GUI contents`). When you open EVE again, the last saved GUI settings will be loaded automatically.
5. **Run box:** When you click run, a full run will be executed.
6. **Preview box:** To check whether the current selection of parameters for the candidate finding is suitable for your data or needs further fine tuning, you can perform a preview run. Doing so, will perform the analysis routines only on a smaller subset of the data that you can specify in the preview box. To view the event data, which in its raw form is just a list of events, it is converted into a format that is easier for humans to view and interpret (images). You must therefore specify a `display frame time` together with the data selection you would like to display. Via `Display event type` you can decide how the event data should be transformed to images You can either preview the finding and fitting results (by pressing `Preview finding/fitting`) or display only the raw events (by pressing `Preview events`).
7. **Tab menu:** Here, you can change between different tabs: `Processing` (current tab), `Post-processing` (follow up analysis after a full run), `Localization List` (view, import and export localization tables), `Visualization` (visualize the super-resolved event SMLM data), `Run Info` (info about current run), `Preview run` (preview of finding and fitting performance, useful for parameter tweaking) and `Candidate Preview` (view single candidate clusters and their x,y,t localization results)

#### 2. Perform a Preview Run

Now, change the `Duration` in the preview box to 10000 ms and then press `Preview`. This will immediately open the run info. By leaving the settings `min` and `max` for x,y empty, you will get a preview for the entire FOV without spatial restrictions.

Switch to the `Preview run` tab, as soon as the preview run is complete (`UpdateShowPreview ran!` is printed in `Run info` tab).

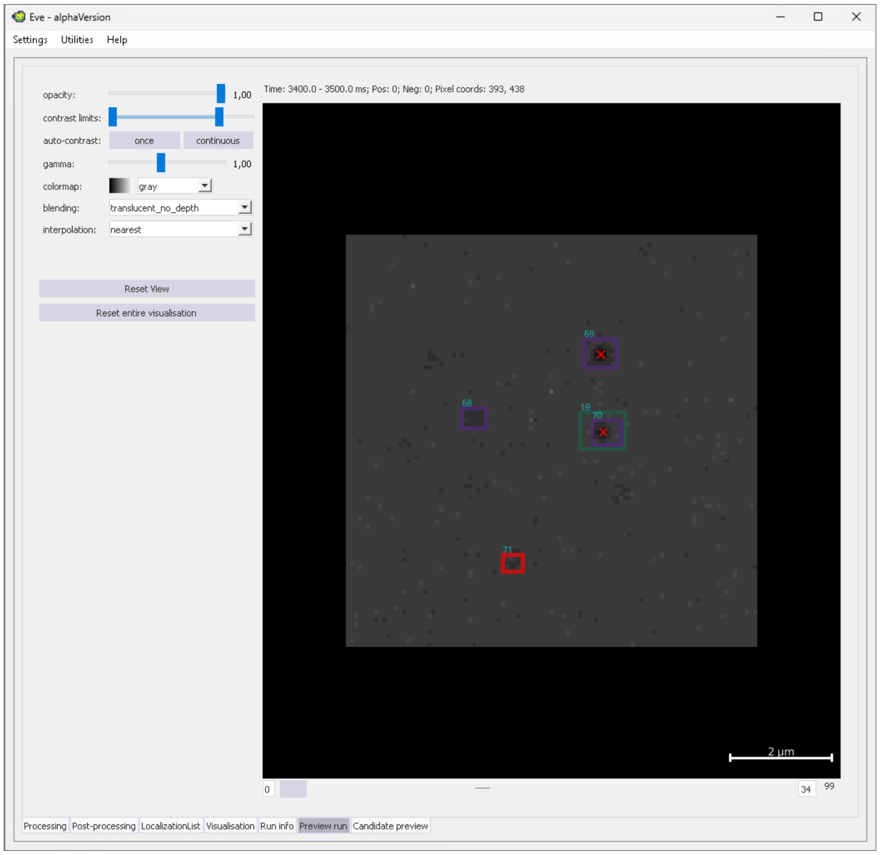


Each candidate cluster found by the candidate finding routine will be highlighted with an purple or green box and gets a unique candidate ID that is here displayed in blue. A red box indicates that a failed fit, meaning that no localization could be generated. All localizations are marked as red crosses in the frame where the x,y,t localization is found. You can use the slider below the image to view all the frames that were created in the preview run.

#### 3. Explore the Candidate Preview

By either double-clicking on a candidate in the preview image or switching to the last tab, you can open the `Candidate preview`.
Change the plot options for the first plot to `3D point cloud of the candidate cluster` and for the second plot to `2D projections of candidate cluster`.
In a preview run you can also display surrounding events to evaluate if the full candidate cluster is found. To do so, set `show surrounding` to `True` and add a custom x,y and t-padding in the first plot options.

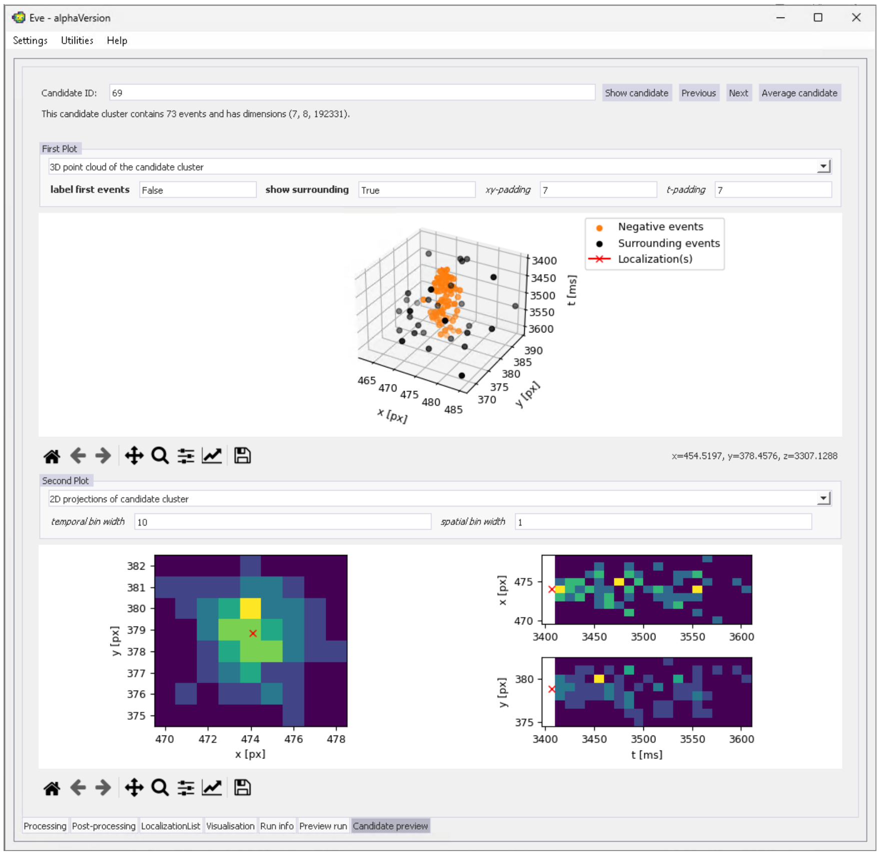


By the `Previous` and `Next` buttons you can simply click through all the clusters found to evaluate finding and fitting results.

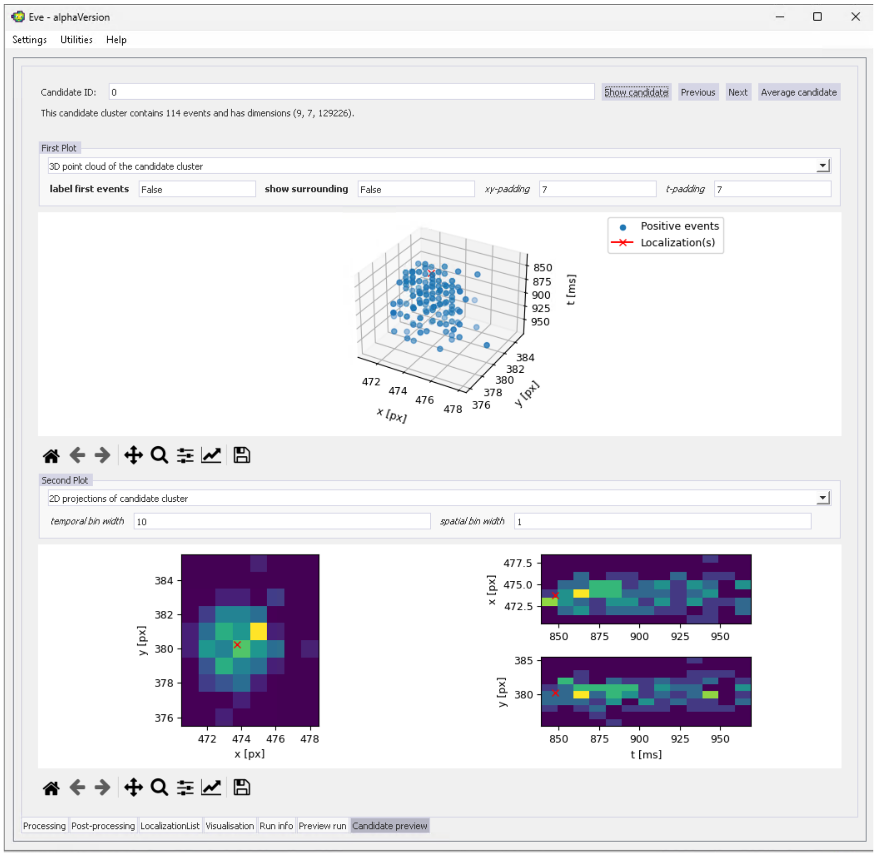

#### 4. Notes of best finding/fitting algorithms and parameters

Knowing which finding/fitting algorithms are best is a currently unsolved problem, and part of why EVE has its expandability. The `Preview run` and `Candidate preview` are designed specifically to give a user the required tools to fully explore the finding and fitting routines. Nonetheless, here are some recommendations to find the best finding and fitting routines at the beginning of a new dataset:

For finding routines, it is recommended to first explore the `Eigen-feature analysis` method via `Preview`, which appears to work robustly. First, set `Maximum Eigenvalue cutoff` to `0`, and set `Debug Boolean` to `True`: Now, a debug figure will be shown. This figure contains a histogram of the maximum Eigenvalue for all events using these settings. Two populations should be discriminatable: Low values correspond to single-molecule clusters, while high values correspond to noise. Close the figure, and change the `Number of neighbours` to higher and lower values (e.g. `10` and `40`), and explore whether this discrimination becomes more pronounced or less pronounced. Do the same with `Ratio ms to px`, and note the single value that most clearly separates the two populations. Next, use this value in the `Maximum Eigenvalue cutoff`, and set `Debug Boolean` to `False`. Re-run the `Preview` and inspect the found clusters. If clusters are missed, try increasing `DBSCAN epsilon` and/or decreasing `DBSCAN nr. neighbours`. If clusters are over-represented, try decreasing `DBSCAN epsilon` and/or increasing `DBSCAN nr. neighbours`.

For fitting routines, it is recommended to start with `2D Logarithmic Gaussian` with `distribution` set to `2D histogram of x,y positions` and `Time fit routine` set to `2D Gaussian (first events)`, if normal, 2-dimensional SMLM is performed. The `expected width` value should be set to roughly the expected sigma of the Point Spread Function (~150 nm for 561 excitation). Then, in `Candidate preview`, loop through a few candidates to see if the found localization agrees with visual inspection, both in its spatial and temporal profile. If many clusters seem to provide a localization where visually not enough events are present, try either decreasing the `fitting tolerance` (which leads to more fits being discarded), or try changing the `Finding` parameters so noisy clusters are discarded. If visually very good clusters are not localized and show a `No localization generated due to ...`-error, try increasing `fitting tolerance`.

#### 5. Execute a full Run

If you are satisfied with the results of the current selection of finding and fitting parameters, you can start a complete run. To do so, switch back to the `Processing` tab and click `Run`.
The `Run Info` tab will again open automatically and show additional info regarding the current run, e.g. number of candidates and valid localizations found as well as a number of candidates (absolute and percentage) that was removed during fitting.

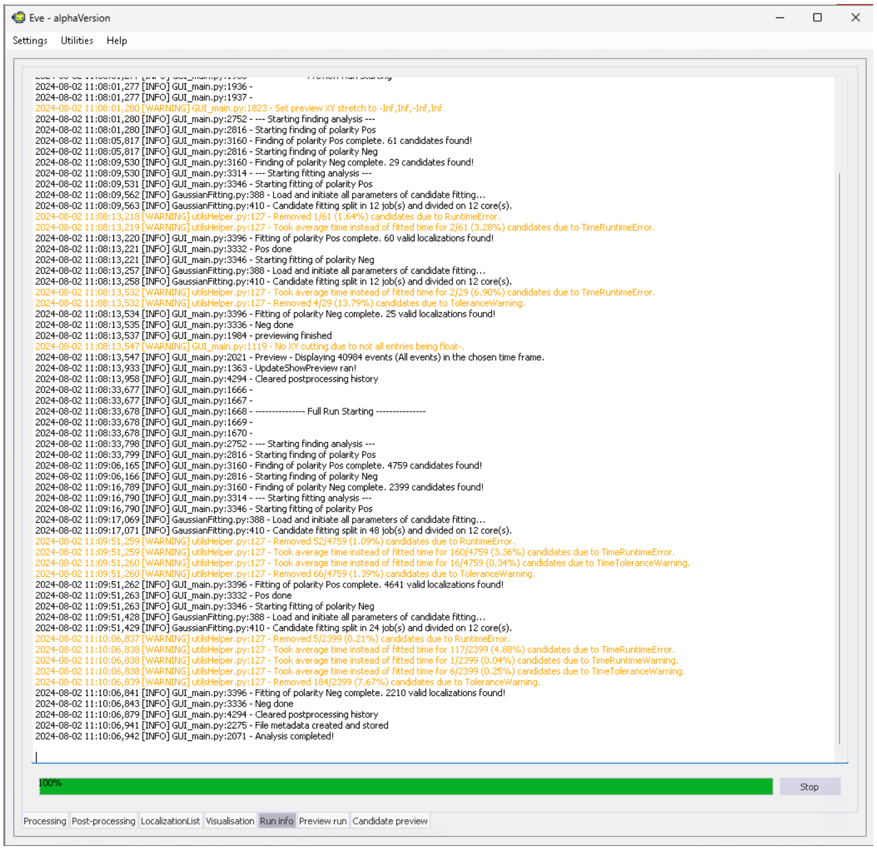


#### 6. Visualize the Localization Results

As soon as the full run is completed, you can visualize your results. Therefore, switch to the `Visualisation` tab, select `2D Histogram with circular kernel convolution` and press `Visualise`.

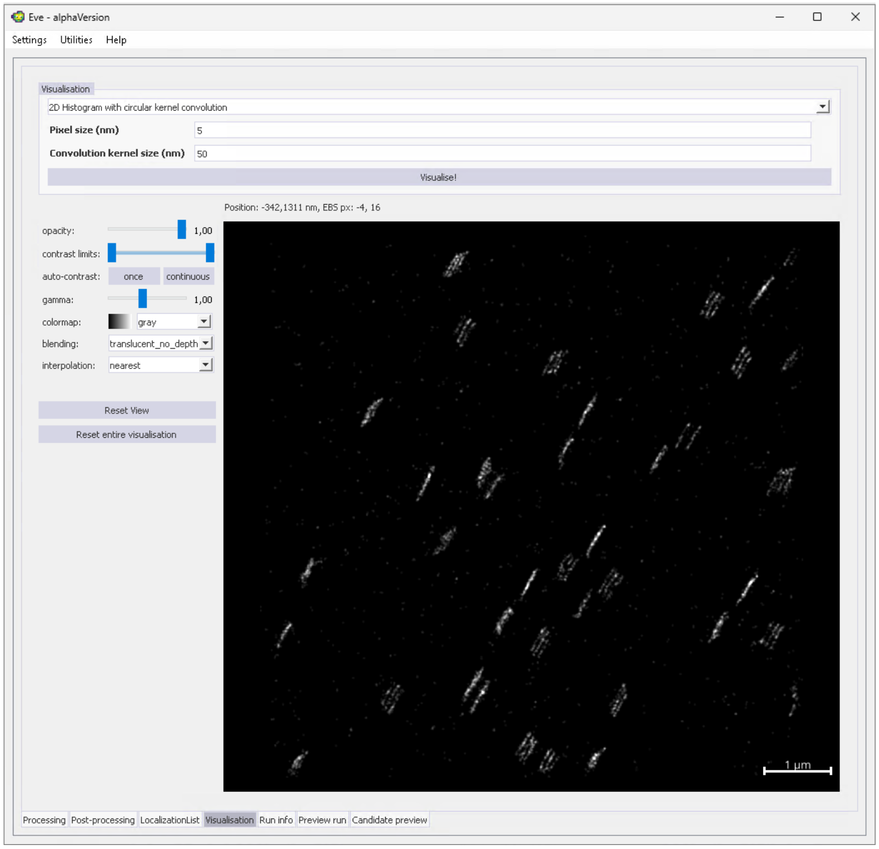


As you can see the sample data is rather drifty and we can't see our DNA nanorulers yet.

#### 7. Apply a Drift Correction

To apply a drift correction, switch to the `Post-processing` tab.

##### Under Windows

Windows users can choose between two different drift correction routines based on entropy minimization or redundant cross-correlation. Select `Drift correction entropy minimization [2D]` and press `Post processing!`. This will open a pop-up window showing the estimated x,y-drift. Additionally, an entry was added to the `Post-processing history` at the bottom of the tab. By clicking `Restore to before this` you can undo the last post-processing step.

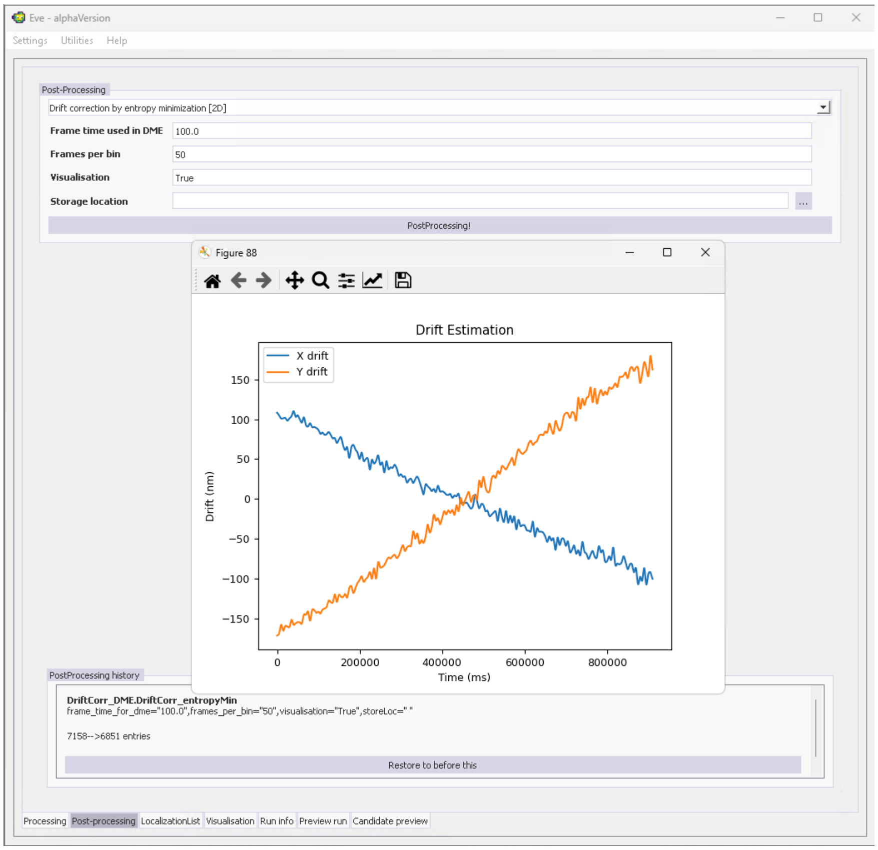


#### Under Linux/MacOs

For Linux and MacOs users currently only one method (`Drift correction by RCC (redundant cross-correlation)`) is working. Select the method and set the `Use ConvHist(Linux/MacOs)` flag to `True`. Now press `Post processing!` which will open a pop-up window showing the estimated x,y-drift. Additionally, an entry was added to the `Post-processing history` at the bottom of the tab. By clicking `Restore to before this` you can undo the last post-processing step.

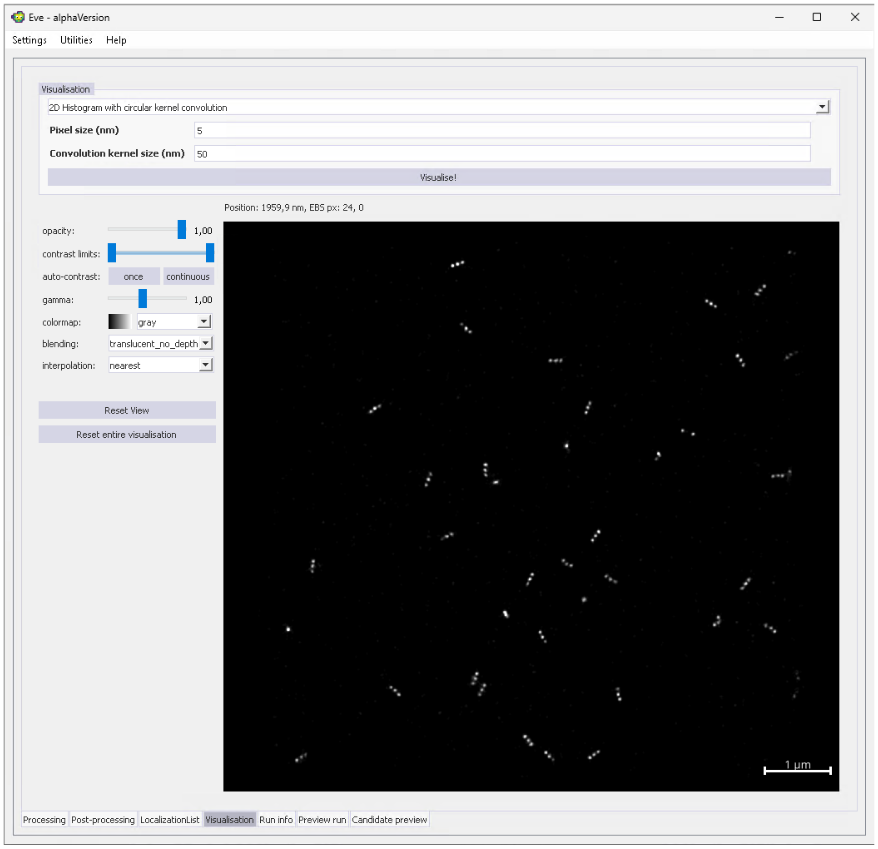


#### 8. The final Localization List

The drift corrected localization list is not saved automatically. To export the list in `.csv` format, switch to the `Localization List` tab and press `Save CSV`.

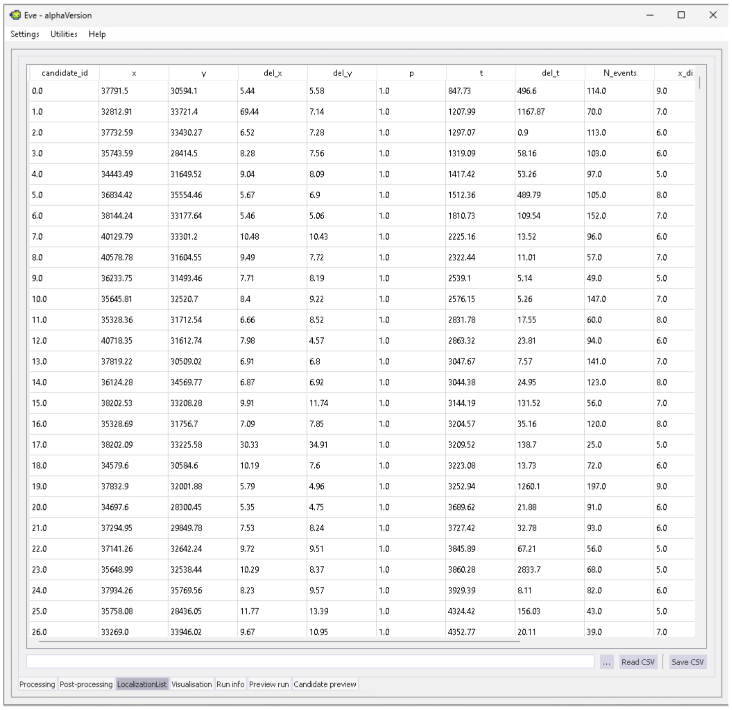

| candidate_id | x | y | del_x | del_y | p | t | del_t | N_events | x_di |
|---|---|---|---|---|---|---|---|---|---|
| 0.0 | 37791.5 | 30594.1 | 5.44 | 5.58 | 1.0 | 847.73 | 496.6 | 114.0 | 9.0 |
| 1.0 | 32812.91 | 33721.4 | 69.44 | 7.14 | 1.0 | 1207.99 | 1167.87 | 70.0 | 7.0 |
| 2.0 | 37732.59 | 33430.27 | 6.52 | 7.28 | 1.0 | 1297.07 | 0.9 | 113.0 | 6.0 |
| 3.0 | 35743.59 | 28414.5 | 8.28 | 7.56 | 1.0 | 1319.09 | 58.16 | 103.0 | 6.0 |
| 4.0 | 34443.49 | 31649.52 | 9.04 | 8.09 | 1.0 | 1417.42 | 53.26 | 97.0 | 5.0 |
| 5.0 | 36834.42 | 35554.46 | 5.67 | 6.9 | 1.0 | 1512.36 | 489.79 | 105.0 | 8.0 |
| 6.0 | 38144.24 | 33177.64 | 5.46 | 5.06 | 1.0 | 1810.73 | 109.54 | 152.0 | 7.0 |
| 7.0 | 40129.79 | 33301.2 | 10.48 | 10.43 | 1.0 | 2225.16 | 13.52 | 96.0 | 6.0 |
| 8.0 | 40578.78 | 31604.55 | 9.49 | 7.72 | 1.0 | 2322.44 | 11.01 | 57.0 | 7.0 |
| 9.0 | 36233.75 | 31493.46 | 7.71 | 8.19 | 1.0 | 2539.1 | 5.14 | 49.0 | 5.0 |
| 10.0 | 35645.81 | 32520.7 | 8.4 | 9.22 | 1.0 | 2576.15 | 5.26 | 147.0 | 7.0 |
| 11.0 | 35328.36 | 31712.54 | 6.66 | 8.52 | 1.0 | 2831.78 | 17.55 | 60.0 | 8.0 |
| 12.0 | 40718.35 | 31612.74 | 7.98 | 4.57 | 1.0 | 2863.32 | 23.81 | 94.0 | 6.0 |
| 13.0 | 37819.22 | 30509.02 | 6.91 | 6.8 | 1.0 | 3047.67 | 7.57 | 141.0 | 7.0 |
| 14.0 | 36124.28 | 34569.77 | 6.87 | 6.92 | 1.0 | 3044.38 | 24.95 | 123.0 | 8.0 |
| 15.0 | 38202.53 | 33208.28 | 9.91 | 11.74 | 1.0 | 3144.19 | 131.52 | 56.0 | 7.0 |
| 16.0 | 35328.69 | 31756.7 | 7.09 | 7.85 | 1.0 | 3204.57 | 35.16 | 120.0 | 8.0 |
| 17.0 | 38202.09 | 33225.58 | 30.33 | 34.91 | 1.0 | 3209.52 | 138.7 | 25.0 | 5.0 |
| 18.0 | 34579.6 | 30584.6 | 10.19 | 7.6 | 1.0 | 3223.08 | 13.73 | 72.0 | 6.0 |
| 19.0 | 37832.9 | 32001.88 | 5.79 | 4.96 | 1.0 | 3252.94 | 1260.1 | 197.0 | 9.0 |
| 20.0 | 34697.6 | 28300.45 | 5.35 | 4.75 | 1.0 | 3689.62 | 21.88 | 91.0 | 6.0 |
| 21.0 | 37294.95 | 29849.78 | 7.53 | 8.24 | 1.0 | 3727.42 | 32.78 | 93.0 | 6.0 |
| 22.0 | 37141.26 | 32642.24 | 9.72 | 9.51 | 1.0 | 3845.89 | 67.21 | 56.0 | 5.0 |
| 23.0 | 35648.99 | 32538.44 | 10.29 | 8.37 | 1.0 | 3860.28 | 2833.7 | 68.0 | 5.0 |
| 24.0 | 37934.26 | 35769.56 | 8.23 | 9.57 | 1.0 | 3929.39 | 8.11 | 82.0 | 6.0 |
| 25.0 | 35758.08 | 28436.05 | 11.77 | 13.39 | 1.0 | 4324.42 | 156.03 | 43.0 | 5.0 |
| 26.0 | 33269.0 | 33946.02 | 9.67 | 10.95 | 1.0 | 4352.77 | 20.11 | 39.0 | 7.0 |

#### 9. Estimate the Localization Precision

To get an estimate on the localzation precision we can make use of the fact, that for each fluorophore blink we measure the on and off-switching. This means, that we have one cluster of positive polarity for the on switching and one cluster of negative cluster polarity for the off switching. By comparing the distances between the corresponding positive and negative localizations we can get our localization precision via a modiefied Nearest neighbour analysis (NeNA).
To do so, we first need to do the match positve and negative localizations that correspond to the same emitter. Therefore, switch again to the `Post-processing` tab and select `Polarity Matching - match on and off events` and press `PostProcessing!` and select appropriate parameters.

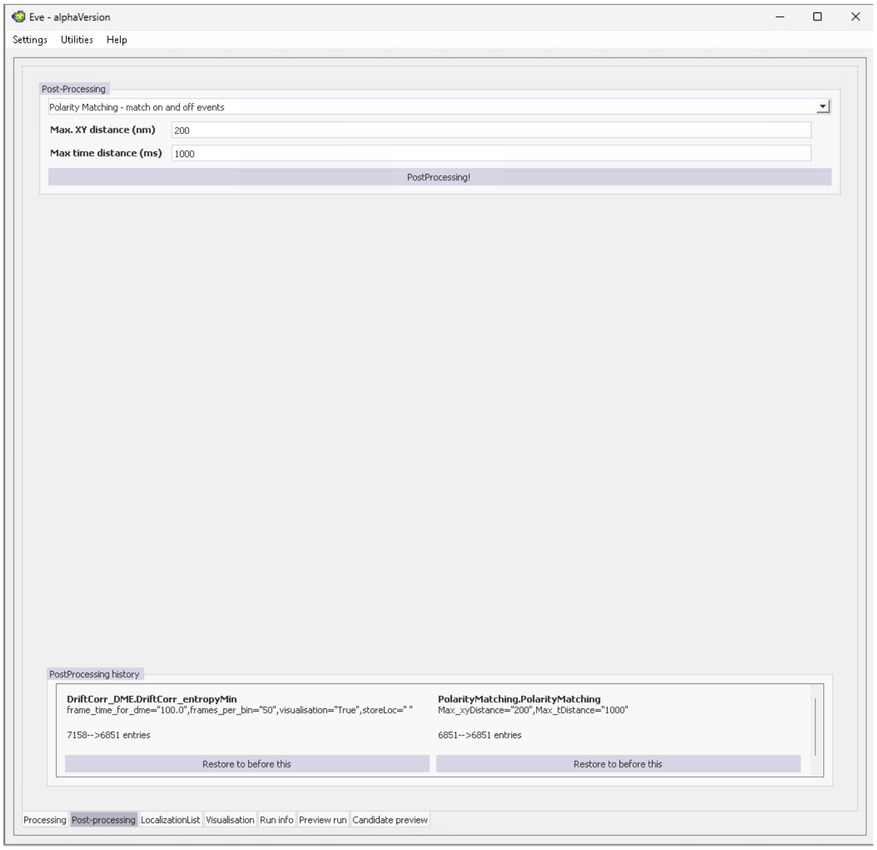


Now, you can perform a NeNA fit by selecting `Nearest neighbour analysis (NeNA) precision on matched polarities` and pressing `PostProcessing!`. This will open a pop-up window showing the distance distribution of all matched localizations along with the NeNA fit. For our DNA nanoruler and the candidate finding and fitting methods and parameters we get a localization precision of ~8.9 nm.

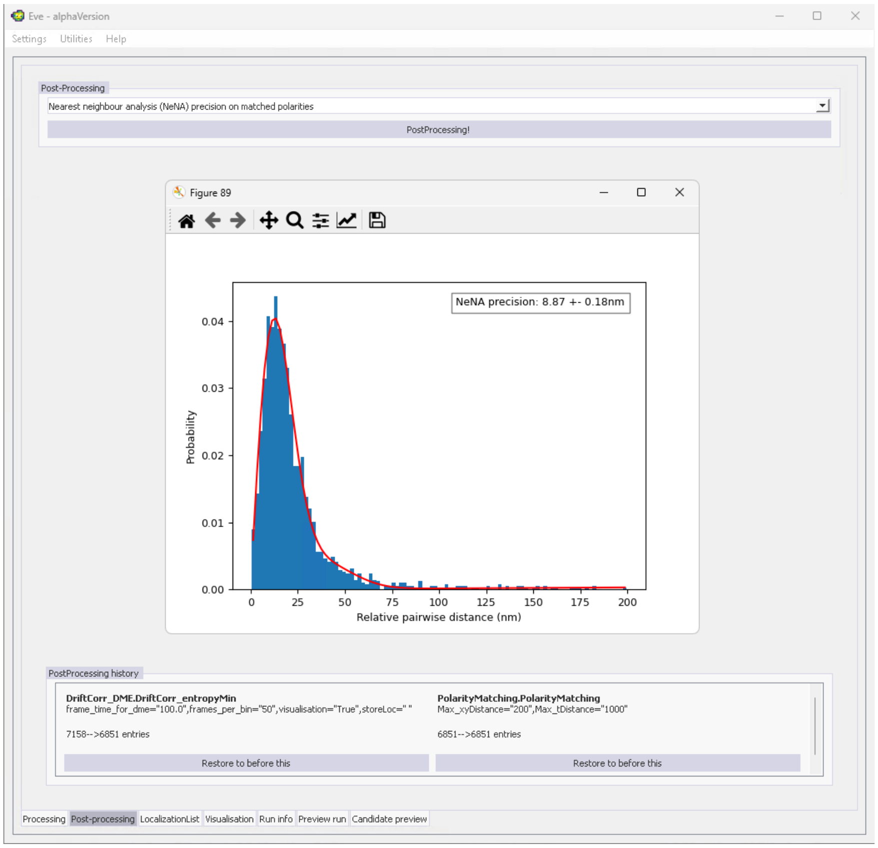


#### 10. Metadata, run and result files

If selected correctly in the `Advanced Settings`, EVE will store a lot of metadata, run and result files automatically. In our "Getting Started" example, three finding and three fitting `.pickle` files are stored containing all, only positive and only negative candidates/localizations. In addition, a `.csv` file with all localizations and a metadata `Runinfo` file are stored.

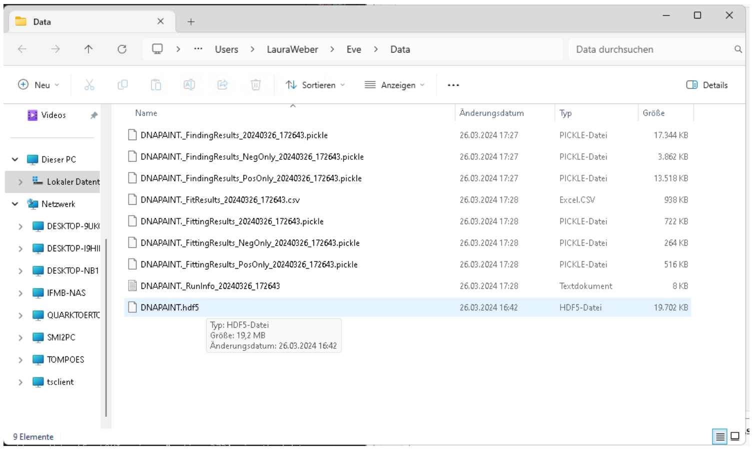


#### 11. Command-line interface

All of EVE can also be accessed via the command-line. For a detailed overview look at `EVE_CommandLine.ipynb`.

#### 12. Exemplary data

The EVE software is packaged with three exemplary datasets, found in the **Data** folder: DNA-PAINT nanoruler, *E. coli* with endogenous RpoC-mEos3.2 and Nile Red membrane stain. The $\alpha$-tubulin labeled Cos7 dSTORM sample dataset, as well as full-sized versions of the other datasets, can be found at doi: 10.5281/zenodo.13269600. Here are succint instructions for how to fully analyse these datasets:

1. Nanoruler (DNA-PAINT):
   - In EVE, in the processing tab, set the Dataset location to the `/Data/Nanoruler/Nanoruler_Small.hdf5` file (via the `File...` button).
   - By using `Settings-Load Specific GUI contents`, load the `/Data/Nanoruler/GUI_Settings_Nanoruler.json` file.
   - Open the advanced settings via `Settings-Advanced Settings`, and use `Load Global Settings` to load the `/Data/Nanoruler/GUI_Settings_Nanoruler_advancedSettings.json` file.
   - Run the analysis by pressing `Run` in the Processing tab, or alternatively, investigate the finding/fitting options via the `Preview` tab.
   - After running, the results need to be drift-corrected - either via `Post-processing - Drift correction by entropy minimization [2D]` (default settings), or by loading (`Load stored drift correction DME/RCC`) the `/Data/Nanoruler/DME_driftCorrection.npz` file.
   - The results can be visualized by opening the `Visualization` tab and running `Gaussian-blurred with locPrecision sigma`.
   - The results should be similar to the localizations found in `/Data/Nanoruler/Nanoruler_Small_FitResults_Drift`[illegible]
2. *E. coli* datasets (PALM and PAINT):
   - Two datasets are provided: `Ecoli_RpoC_Small` and `Ecoli_NR_Large`, along with their corresponding GUI settings. Throughout these instructions, these can be interchanged to obtain the results for either dataset.
   - In EVE, in the processing tab, set the Dataset location to the `/Data/Ecoli/Ecoli_NR_Small.hdf5` file (via the `File...` button).
   - By using `Settings-Load Specific GUI contents`, load the `/Data/Ecoli/GUI_Settings_EColi_NR.json` file.
   - Open the advanced settings via `Settings-Advanced Settings`, and use `Load Global Settings` to load the `/Data/Ecoli/GUI_Settings_EColi_NR_advancedSettings.json` file.
   - Run the analysis by pressing `Run` in the Processing tab, or alternatively, investigate the finding/fitting options via the `Preview` tab.
   - The results can be visualized by opening the `Visualization` tab and running `Gaussian-blurred with locPrecision sigma`.
   - These results should be similar to the localizations found in `/Data/EColi/NEcoli_NR_Small_FitResults.csv`.
3. *α-tubulin* in Cos-7 cell (dSTORM):
   - Download the dataset from doi:10.5281/zenodo.13269600.
   - In EVE, in the processing tab, set the Dataset location to the `/Data/aTubulin/aTubulin_Small.hdf5` file (via the `File...` button).
   - By using `Settings-Load Specific GUI contents`, load the `/Data/aTubulin/GUI_Settings_aTubulin.json` file.
   - Open the advanced settings via `Settings-Advanced Settings`, and use `Load Global Settings` to load the `/Data/aTubulin/GUI_Settings_aTubulin_advancedSettings.json` file.
   - Run the analysis by pressing `Run` in the Processing tab, or alternatively, investigate the finding/fitting options via the `Preview` tab.
   - After running, the results need to be drift-corrected - either via `Post-processing - Drift correction by entropy minimization [2D]` (default settings), or by loading (`Load stored drift correction DME/RCC`) the `/Data/aTubulin/DME_driftCorrection.npz` file.
   - The results can be visualized by opening the `Visualization` tab and running `Gaussian-blurred with locPrecision sigma`.
   - These results should be similar to the localizations found in `/Data/aTubulin/aTubulin_Small_FitResults.csv`.

### Supplementary Note 3: Developer Instructions - EVE software

#### Contents


- Introduction
- Expandability of EVE
- Detailed information on input/output data of EVE
  - Candidate Finding
  - Candidate Fitting
  - Post-processing
  - Visualization
  - Candidate preview
  - Event distributions
  - Temporal Fitting

### Introduction

EVE is a software platform developed for the analysis of single-molecule imaging data captured by event-based sensors. This document is for developers who want to add functionality to EVE. EVE is a highly open framework and can easily be expanded upon. Expandability of EVE is possible in the following routines (with more detailed information following):

- **Candidate Finding**

  Routines involved in finding which events belong to a single molecule localization/PSF

  **Input**: All events (possibly filtered by polarity), settings, function arguments

  **Output**: Found candidates, metadata

- **Candidate Fitting**

  Routines involved in fitting the events of each candidate cluster to determine the x-, y- (,z-) and t-coordinates of each single molecule. Some fitting methods rely on using 'Event distributions' (below)

  **Input**: All found candidates, settings, function arguments

  **Output**: Localizations (x,y(,z),t-coordinates), metadata

- **Event distributions**

  Classes to create varying distributions from events

  **Input**: Events, settings, function arguments

  **Output**: Histogram classes with certain arguments

- **Post-processing**

  Routines involved in post-processing the localization data, either for further filtering or data quantification or for calculating quantitative metrics

  **Input**: Localizations, candidates, settings, function arguments

  **Output**: (Possibly changed) localizations, metadata; or None

- **Visualization**

  Routines that visualize the current localization list

  **Input**: Localizations, settings, function arguments

  **Output**: 2D array containing image data, scale of the image

- **Candidate preview**

  Routines that visualize individual candidates for user inspection

  **Input**: Candidates, localizations, all events, settings, function arguments

  **Output**: None (updated figure)

### Expandability of EVE

All routines, with the exception of the Event distributions, follow the same method of expandability. One or multiple routines should be written in a .py file, and placed within a sub-folder in the main EVE GUI folder, or alternatively in the AppData/Local/UniBonn/Eve folder. EVE will automatically find and add all suitable routines into the GUI. The .py files should start with the following defined structure:

```
def __function_metadata__():
 return {
  "FunctionTitle": {
   "required_kwargs": [
    {"name": "kwarg1", "description": "Some Description", "default": 5.0, " type": float,
    "display_text": "Keyword Argument 1"},
    {"name": "kwarg2", "description": "Some other Description", "default":"string",
    "type": str, "display_text": "Keyword Argument 2"}
   ],
   "optional_kwargs": [
    {"name": "oarg", "description": "An optional argument", "default":1, "type":int,
    "display_text":"Optional Argument"},
   ],
   "help_string": "This is an exemplary routine.",
    "display_name": "Exemplary Routine"
  }
 }
```

This function would be displayed as such by the EVE GUI (exemplary for candidate finding on positive events):

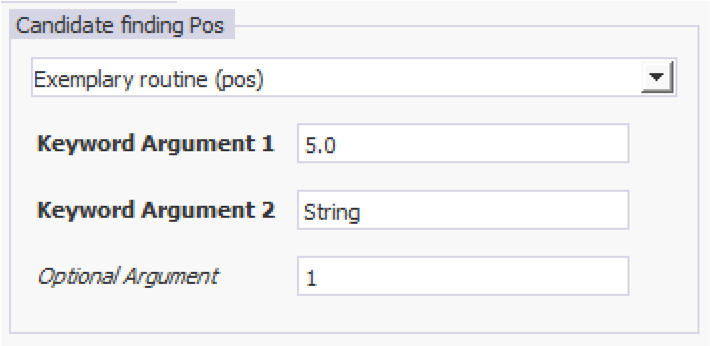


The function_metadata function describes important metadata for the function(s) that should be displayed and callable:

- One or multiple functions can be defined with this structure
- The same .py file requires a function called the same as “FunctionTitle”

The following parameters should be created:

**display__name** (optional) provides the function name which is visible for the user

**help__string** (optional) provides a description of the function for the user

**required__kwargs** (required)** defines required keyword arguments that your function expects.

**optional__kwargs** (required)** defines optional keyword arguments that your function could use.

- For each keyword argument, the following should be provided:

  name (required): the internal name of the keyword argument

  display_text (optional): the name visible to the user

  description (optional): a description of the argument that users can see in the EVE GUI default (optional): the default value of this argument

  type (optional): the expected type of the input, options are [float, int, str, "fileLoc"]

  The "fileLoc" value indicates a file which can be found by the user

### Detailed information on input/output data of EVE

All data has these two input variables:

*settings*: named dictionary with (advanced) settings.
*kwargs*: dictionary with named entries of the function (as defined in *__function_metadata__()*)

#### Candidate Finding

**Function definition**
*def function(npy_array, settings,**kwargs): return candidates, performance_metadata*

**Input**
*npy_array*: numpy.ndarray with one entry for each event. Each entry has dtype([(‘x’, ‘<u2’), (‘y’, ‘<u2’), (‘p’, ‘<i2’), (‘t’, ‘<i8’)]) structure, with x/y in pixels, p either 0 or 1 (negative or positive), and t in microseconds

**Output**
*candidates*: dictionary where each entry is a candidate. Each entry should have three named sub-entries:

*events:* pandas DataFrame with N-by-4 array, array names x,y,t,p (same units as input, N being the number of events in this cluster).

*N_events*: Number of events

*cluster_size*: [size_x, size_y, size_t] of the cluster (in [pixel, pixel, microsecond] units)

*performance_metadata*: string with details on the performance. Will be stored in the metadata.txt output.

##### Pseudo-code explaining the structure of candidates finding output

```
candidates = {}
for cluster in all_clusters:
    clusterEvents = all_cluster_events(cluster_id==cluster)
    candidates[cluster] = {}
    candidates[cluster]['events'] = clusterEvents
    candidates[cluster]['N_events'] = len(clusterEvents)
    candidates[cluster]['cluster_size'] =...
    [np.max(clusterEvents['y'])-np.min(clusterEvents['y']),...
    np.max(clusterEvents['x'])-np.min(clusterEvents['x']),...
    np.max(clusterEvents['t'])-np.min(clusterEvents['t'])]

metadata = 'The file ran as expected!'
```

### Candidate Fitting

For the candidate fitting, the *__function_metadata__()* needs to be expanded to provide information about the dist_kwarg and time_kwarg. These structures contain information about which XY distribution and Time distribution can be selected by the user. If these are not defined, an XYT-combined fitting is ran (which should result in XY ánd time fitting results). In an XY+Time distribution, the candidate fitting routine should only provide the XY fitting result, since the Time distribution is handled independently. Please look at the following examples for implementation details:

Example for XY+Time: GaussianFitting

Example for XYT: Radial_Symmetry – RadialSym3D.

**Function definition** *def function(candidate_dic, settings,**kwargs): return localizations, fit_info

**Input** *candidate_dic: see output from candidate finding

**Output** *localizations*: Pandas Dataframe of localizations corresponding to input clusters. *fit_info*: string with info of metadata. Will be stored in the metadata.txt output. Should at least have columns with names 'candidate_id', 'x', 'y', 'p', ['t'], in units integer, pixel, pixel, 0/1, microseconds, respectively. t is not required if an independent Time distribution fitting is used (see above). Can also have more columns as wanted, with nomenclature normally following 'del_x' for uncertainty in x. Commonly also 'fit_info' column can be used to report on incomplete/wrong fits. Each candidate should have one entry. *performance_metadata*: string with details on the performance. Will be stored in the metadata.txt output.

#### Pseudo-code explaining the structure of candidate fitting output

```
localizations = {}
for i in np.unique(list(candidate_dic)):
    localizations[i]={}
    localizations[i]['x'] =
    np.mean(candidate_dic[i]['events']['x'])*float(settings['PixelSize_nm']['value']) #X position in nm
    localizations[i]['y'] =
    np.mean(candidate_dic[i]['events']['y'])*float(settings['PixelSize_nm']['value']) #Y position in nm
    localizations[i]['p'] = 1 #Polarisation: 0 or 1
    localizations[i]['t'] = np.mean(candidate_dic[i]['events']['t'])/1000 #time in ms

#Make a pd dataframe out of it - needs to be transposed
localizations = pd.DataFrame(localizations).T

metadata = 'The file ran as expected!'
```

### Post-processing

**Function definition** *def function(localizations, findingResult, settings, **kwargs): return localizations, metadata*

**Input** *localizations*: See Candidate Fitting 'localizations' output. *findingResult*: See Candidate Finding 'candidates' output.

**Output** *localizations*: See Candidate Fitting 'localizations' output. Practically, should be a filtered/amended list of the original localizations. *metadata*: string with info of metadata. Will be shown in the Run info GUI tab.

### Visualization

**Function definition** *def function(resultArray, settings,**kwargs): return image, scale*

**Input** *resultArray*: See Candidate Fitting 'localizations' output. Uses the currently found results as in Eve (i.e. could be adapted via Post-processing).

**Output** *image*: numpy.ndarray of pixel-values of the resulted image. Will be displayed in the 'Visualization' tab *scale*: float value of pixel-to-micrometer size (e.g. value of 0.01 means 0.01 micrometer per pixel, or 10 nm per pixel). Used to set the scale bar in the 'Visualization' tab.

### Candidate preview

**Function definition** *def function(findingResult, fittingResult, previewEvents, figure, settings,**kwargs): return None*

**Input** *findingResult*: See Candidate Finding 'candidates' output. The information of a single candidate is provided. *fittingResult:* See Candidate Fitting 'localizations' output. The information of a single localization is provided. *previewEvents*: Unused *figure*: Matplotlib Figure object. Should be addressed by e.g. performing *ax = figure.add_subplot(111); ax.bar(...).* figure.show() does not have to be called.

**Output** *None*. Expected that *figure* is updated properly.

### Event distributions

These Event distributions follow a different expandability method, and cannot be adapted from the AppData folder, but only from changing the EventDistributions/eventDistributions.py file in the EVE installation folder.

Each Event distribution is defined by a class (e.g. *class Hist1d_t()* ). These classes should have a *___call___(self, events, **kwargs)* function, which should return the wanted distribution and bin edge positions.

Please use the existing classes in *eventDistributions.py* for detailed info.

### Temporal Fitting

Same structure as Event distributions, only for fitting time distributions.
